# Gene transcription and chromatin packing domains form a self- organizing system

**DOI:** 10.64898/2026.03.15.711889

**Authors:** Lucas M Carter, Wing Shun Li, Ruyi Gong, Nicolas Acosta, Niyati Pandya, Marcelo A Carignano, Emily Pujadas Liwag, Kunlun Wang, Tiffany Kuo, Kyle MacQuarrie, Masato T. Kanemaki, Luay Almassalha, Vadim Backman

## Abstract

The human genome organizes into several thousand chromatin packing domains that couple euchromatin and heterochromatin into unified nanoscale volumes. Prior work suggested these domains form by transcriptionally driven loops that guide packing domain assembly. Here, we study the process of transcriptionally driven domain formation and maintenance. By pairing auxin-inducible degron technology with nanoscopic imaging, transcriptomics, and Hi-C, we show that Pol-II regulates conformationally defined interphase chromatin packing domains. Pol-II facilitates nascent domain generation and maintains mature domain integrity through the process of generating transcriptional loops. Mechanistically, Pol-II maintains the packing of intronic and intergenic chromatin within domains by transcribing exons within gene bodies. Consequently, polymerase loss disrupts genome connectivity, *in situ* packing domains, and gene expression, genome-wide. Our findings suggest chromatin packing domains and RNA synthesis are tightly coupled to optimize transcriptional responses in human cells.

## Introduction

The irregular 10-nm “beads-on-a-string” chromatin heteropolymer is the fundamental unit of the mammalian genome. Above this scale, 3D chromatin structure is an area of active investigation, with many distinct connectivity and structural features described in literature. There are two complementary approaches to study the structure-function of the genome at these scales: measures of genome connectivity (Hi-C, GAM, Chia-PET, etc.) and measures of physical geometry (SMLM, ChromEM, ORCA, etc). Topologically associated domains (TADs), chromatin loops, and A/B compartments are widely observed across species, indicative of a robust genome connectivity framework^1,2^. Connectivity assays are limited by the inability to capture physical packing geometry, while conversely, imaging often lacks sequence-specific regulatory information. As a result, the mechanisms linking topology, geometry, and function within the genome remain poorly defined. To date, relating chromatin states across these modalities was largely built on prior electron microscopy imaging observations which identified two discrete states in genome organization: densely packed heterochromatin (B-compartments, nuclear periphery, transcriptionally inactive) and poorly packed euchromatin (A- compartments, nuclear interior, transcriptionally active)^3,4^.

Recent advances in chromatin electron microscopy (ChromEM) are challenging this underlying binary framework^5,6^. While ChromEM lacks sequence-specific information, signal intensity is proportional to local chromatin density. A key observation from ChromEM is that the fundamental structural unit above the level of individual nucleosomes is a continuous meshwork of supranucleosomal packing domains (PDs). Every PD exhibits a power-law scaling behavior, consistent with fractal packing across length scales. This scaling results in a local density gradient from a high-density core (heterochromatin) to a lower density outer shell (euchromatin). Packing domain geometry couples heterochromatin and euchromatin together, forming an ideal intermediate density interface that optimizes physiochemical conditions for efficient transcriptional reactions near the domain surface. Since PDs have a continuous distribution of sizes (50-300nm) and packing efficiencies (density), prior work proposed that this distribution may relate to a dynamic structure-function lifecycle closely linking transcriptional reactions with physical geometry^6–9^.

Specifically, it has been proposed that active transcription may generate local density gradients wherein the elongating polymerase generates forced returns in the confined space of the nucleus—reconfiguring the chromatin heteropolymer into volumes (domains). An initially weak (nascent) density gradient under sustained transcriptional use biases the movement of chromatin remodeling complexes, such as heterochromatin and euchromatin modifying enzymes, by their size resulting in a process of PD maturation (efficient packing). As a consequence of this maturation, the interface of the PD is stabilized into an “ideal” zone, enhancing future transcriptional reactions. In this framework, transcriptional activity and physical state mutually reenforce each other, giving rise to an emergent structure-function relationship at the nanoscale^7,8^.

Previous investigation relied on complex pharmacological disruption of this structure-function relationship^10,11^, leaving a major question unresolved: does Pol-II directly facilitate the domain lifecycle? Using auxin-inducible degron technology targeting the largest subunit of Pol-II^12,13^, we compared the role of Pol-II in regulating domain contact scaling with Hi-C alongside the *in situ* physical structure of packing domains using the recently developed Nanoscale Chromatin Imaging and Analysis (nano-ChIA) platform that combines chromatin electron scanning transmission microscopy (ChromSTEM), partial wave spectroscopy (PWS), and multiplexed single molecule localization microscopy (SMLM)^14^. Finally, to probe the relationship between structure and function directly, we investigated the loss of Pol-II on gene function using nascent (EU-seq) and total expression analysis (RNA-seq, RIP-seq). Here, we show that Pol-II guides nascent domain formation and mature domain maintenance through transcription. Mechanistically, our results suggest that this process occurs through packing non-transcribed elements, such as introns, into 3D volumes. Consequently, from a physical genomics perspective, we describe how polymerase loss disrupts genome connectivity, *in situ* packing domains, and gene expression genome-wide by disrupting the domain lifecycle.

These observations provide a physical mechanism that may explain why transcription occurs in B compartments and why A compartments are not completely devoid of heterochromatin markers^15,16^. In addition, they may also provide insight into disagreement between observations from different methods used to probe the relationship between 3D structure and transcription. For example, independently degrading all three polymerases produced minimal changes in TADs or compartments in high-throughput chromatin conformation capture (Hi-C)^17^. Likewise, global degradation of cohesin or CTCF, the regulators of TADs, does not lead to wide-scale disruption of transcription^18,19^, but downstream nascent transcription is perturbed in the absence of either^20^. Conversely, pharmacological disruption of transcription profoundly impacts 3D chromatin structure in super resolution imaging studies^10,11,13^. While future work is needed to understand the full mechanistic implications of these observations, they suggest that polymerase function drives the formation and maintenance of 3D chromatin packing structure.

## Results

### The physical framework of the packing domain structure-function lifecycle

The physical properties of chromatin PDs include fractal dimension (*D*), density (chromatin volume concentration, CVC), and domain radius (r). Packing domains *in situ* fold based on the polymeric power-law properties of the genome. These can be quantified by how mass scales with the space the chromatin polymer occupies, *M* ∝ *r*^*D*^, where *D* is the scaling coefficient. In this sense, *D* represents the conformational packing of domains, where low *D* conformations fill space less efficiently and high *D* conformations fill space more efficiently. Inversely proportional to *D* within a PD is the contact scaling exponent, *S*, which defines the contact probability between units as a function of length. For individual cells, measurements between *S* and *D* are inversely related, where 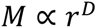, with biologically relevant *D* typically observed between 2 and 3^6,14^. While *S* is typically measured from ensemble Hi-C datasets, these population-averaged measurements may obscure local scaling behaviors present within individual domains^14^.

Indeed, topologic constraints within a population are not necessarily equivalent to individual domains. Crucially, multiple lines of evidence suggest that chromatin PDs are not a physical manifestation of TADs: they are (1) largely stable following RAD21 depletion^21^, (2) directly regulate gene transcription, and (3) depend on polymeric folding in a constrained nuclear volume. Conversely, TADs depend on RAD21 and CTCF for function, are not necessary to maintain transcription, and are not impacted by changes in nuclear volume^18,22^. Given these distinctions, it is necessary to consider how to relate the geometric properties of PDs with the connectivity features in a population ensemble measured by Hi-C.

To establish a framework for comparison, we briefly discuss observations from chromatin polymer modeling that quantitatively and qualitatively recapture the physical properties of PDs. A chromatin polymer without processes that create forced returns (e.g. a self-avoiding walk) adopts a relaxed configuration lacking the domain structures observed via ChromSTEM (**Fig. 1a**). Conversely, introducing stochastically generated forced returns (spatial confinement of segments) is sufficient to generate PDs observed in ChromSTEM experiments (**Fig. 1b**). This observation motivates the hypothesis that enzymatic processes capable of inducing forced returns, such as RNA polymerases, cohesin, or topoisomerases, may facilitate PD formation in a confined environment. While a subset of transcriptionally-mediated forced returns may appear in ensemble Hi-C as enhancer-promoter and promoter-promoter loops ^23^, the stochastic nature of transcription suggests many forced returns may remain undetectable by population-averaged methods^24^. To test the hypothesis that transcription is central to the PD structure-function lifecycle, we first examined the effects of global transcriptional inhibition (via ActD treatment) followed by targeted degradation of RNA polymerase II using the AID2 system^12^.

**Figure 1.**
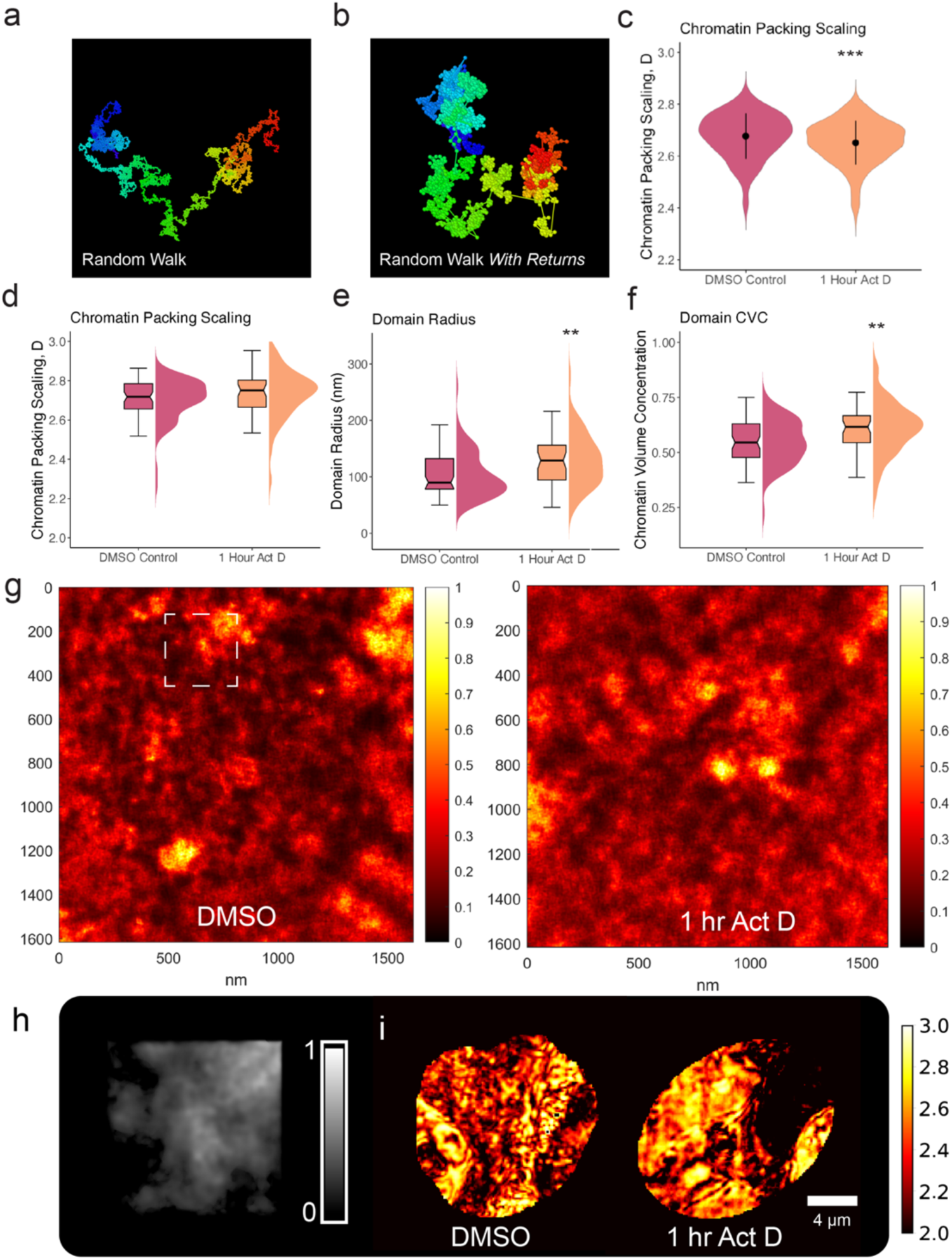
Packing domain analysis following transcriptional inhibition of HCT116 cells treated with Actinomycin D (5 µg/mL) or treated with DMSO. Representative chromatin polymer chains from (**A**) a Self-Avoiding Walk model demonstrating a random walk with no forced returns (enzymatically driven loops) that does not form packing domains and (**B**) a Self-Returning-Excluded Volume model SR-EV demonstrating the formation of packing domains due to the contribution of loops and the excluded volume effect of monomers (nucleosomes). Color corresponds to monomer number in chain: Blue = 1, Red = 4000. (**C**) PWS following ActD treatment: average nuclear packing scaling of packing domains. Data was compiled from three independent biological replicates. (DMSO: N = 475; ActD: N=541). (**D-F**) ChromSTEM following treatment: packing scaling, domain radius, and chromatin volume concentration of individual domains, respectively (DMSO: N = 71; ActD: N=48). (**G**) ChromSTEM tomograms showing untreated chromatin state or domain swelling following treatment. XY axes are in nm. (**H**) Representative domain from ChromSTEM tomogram demonstrating radial density from center to periphery where values approaching 1 have high CVC and values approaching 0 have low CVC (**I**) Representative map of chromatin scaling values, *D*_*nucleus*_, generated using PWS from wildtype nuclei (Left) and ActD-treated nuclei (Right). For **C–F**: Significance was calculated by unpaired *t* test with Welch’s correction applied (**** < 0.0001).

### Inhibiting transcription globally disrupts in situ domains

To evaluate the consequence of inhibiting transcription on domain stability, we began by disrupting transcription of all three polymerases with Actinomycin D (ActD)^7^. While ActD is a well-defined transcriptional disrupter of all three polymerases with dose-dependent selective inhibition of Pol-I, Pol-II, and Pol-III^25–27^, it has a complex mechanism of action with many off-target effects^28–30^. These off-target effects limit the phenotypic interpretation of ActD-treated cells. With this in mind, we conservatively interpreted our results. We used 1 hour ActD treatment (at 5 µg/mL) ^31,32^ to inhibit the activity of all three RNA polymerases and confirmed the arrest of RNA synthesis by 5-Ethynyl-uridine at this concentration (**SI Fig. 1a-c**). To measure the impact on PD structure, we used two components of the Nano-ChIA platform: live-cell Partial Wave Spectroscopy (PWS) nanoscopy and ChromSTEM tomography^14^. PWS is a label-free live-cell imaging technique that measures the interference in backscattered light from chromatin, resolving structures between 20-200 nm^33^. While PWS does not resolve each individual domain structure, it provides information about the distribution of nanoscale packing domain sizes, average domain packing scaling across the nucleus, *D_nucleus_* , domain mobility in live cells, and fractional moving mass (FMM), a measure of coherent polynucleosome movement. For a full discussion of PWS, see Methods. In as little as 1 hour after ActD-mediated inhibition we observed an acute decrease in packing scaling, *D_nucleus_*, and FMM, consistent with the decreased likelihood of chromatin organizing into PDs throughout the nucleus in HCT-116 cells (**Fig. 1c, i**)(**SI Fig. 1d**). To confirm that this is a generalized phenomenon across cell models, we performed similar experiments in three additional cell lines with comparable results (**SI Fig. 1h**).

To understand the effect of inhibiting transcription on individual domains, we performed ChromSTEM tomography (a variant of ChromEM) on cells treated with ActD and on untreated controls. Although ChromSTEM lacks sequence specificity, the resolved structures on ChromSTEM represent the geometric ground truth of the genome *in situ,* where the STEM HAADF contrast intensity is directly proportional to chromatin density with resolution less than 4 nm (**Fig. 1h**). This facilitates the capacity to study the scaling of individual chromatin packing domains, *D*_*domain*_ , whereas PWS can be used to quantify average scaling across the nucleus, *D*_*nucleus*_. On ChromSTEM, we observed that ActD treatment results in a loss of PDs overall, with the primary effect on nascent domains. For the remaining domains, the size, density, and fractal dimension all increased (+26.15 nm, +0.064 CVC, +0.028 D; p < .0001) on average (**Fig. 1d-f**). These changes were also visually apparent on the tomogram generated from our analysis (**Fig. 1g**). Taken together, inhibiting transcription of all three polymerases using Act D produced an aggregation of chromatin into fewer but denser domains, consistent with the hypothesis that transcription regulates the 3D packing of domains. Notably, these findings were consistent with similar observations using super-resolution microscopy in which ActD generates an abrupt DNA-compaction phenotype across the nucleus^10,11^.

### RNA Polymerase II depletion attenuates mRNA synthesis

To overcome the limitations of Act D treatment as a tool for analyzing the contributions of transcription to the PD lifecycle, we utilized a degron line in HCT116 cells targeting POLR2A^12^. For this study, we used an NTD-tagged Pol-II line to ensure efficient degradation upon auxin treatment^34,35^. We first validated the depletion using western blot analysis, wide-field imaging, and flow cytometry, confirming the complete degradation of POLR2A at 6 hours (**SI Fig. 1e-g**). Next, we used EU staining to quantify the effect of depletion at 6 hours on RNA synthesis. Surprisingly, although POLR2A depletion attenuated transcription in the nucleoplasm, limited RNA synthesis was still apparent *in situ*. This was observed both on widefield imaging (**Fig. 2a-b**) and on single molecule localization microscopy (SMLM, **Fig. 2c-g**). To verify that this unexpected result was not due to the diffusion of rRNA from the nucleolus or alternative RNA transcripts, we performed total RNA-Seq on ActD-treated cells (positive control), POLR2A depleted cells (1 µM 5-ph-IAA, 8 hours) and wildtype HCT116 cells (DMSO, 8 hours). Using a liberal cutoff (Padj. < 0.05, abs(LFC) > 1), we observed broad suppression of gene transcription with ActD (**Fig. 2h**).

**Figure 2.**
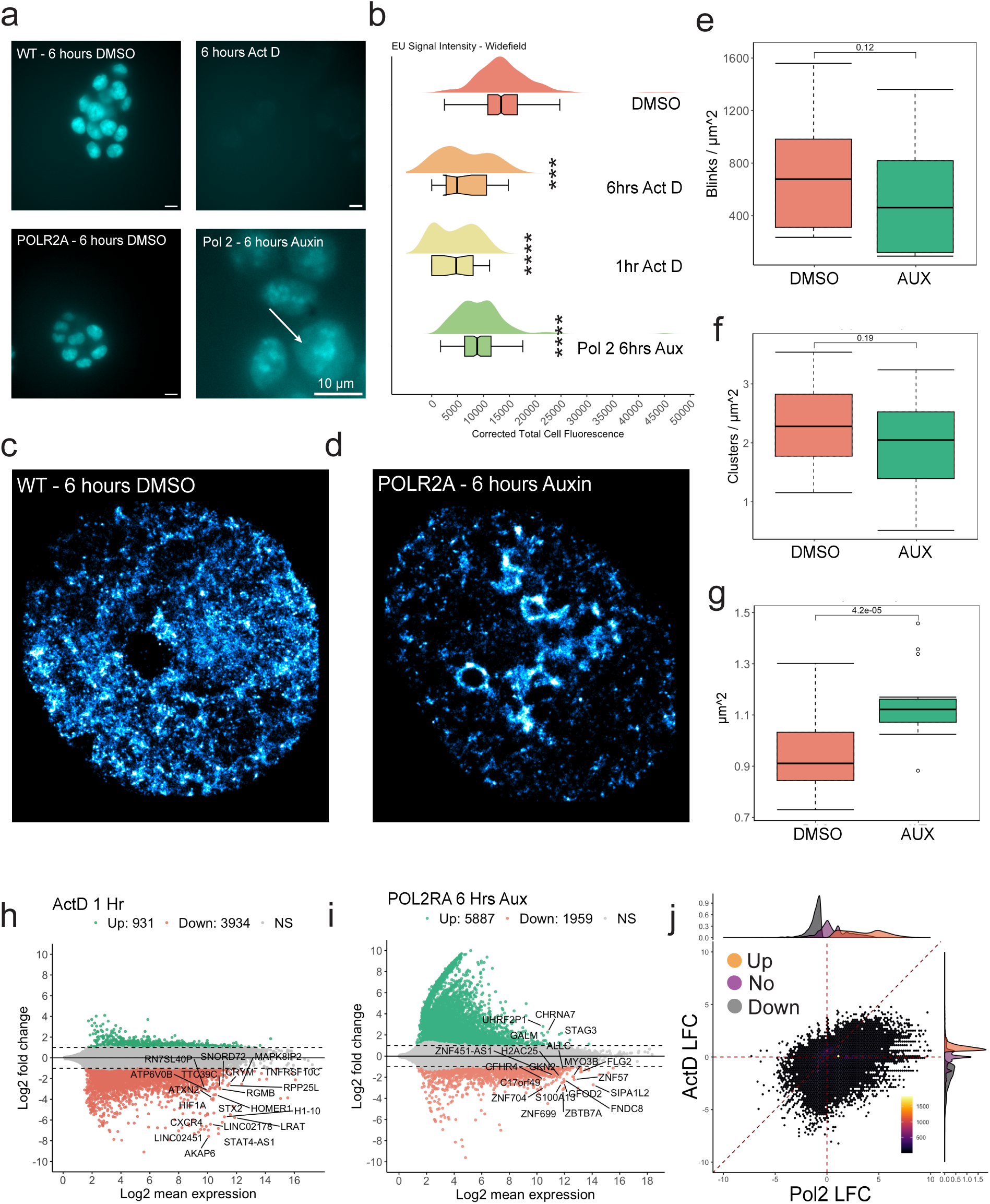
RNA synthesis is downregulated but gene transcription persists in nucleoplasm despite Pol-II depletion. POLR2A-AID2 Degron line was treated with auxin, DMSO, or Actinomycin D treated (5 µg/mL) for 6 hours and imaged following labeling of nascent RNA with 1 mM 5-ethynyl-uridine for 1 hour. (**A**) Representative image of each condition captured using widefield fluorescent microscopy. Scale bar = 10 µm. (**B**) Cell total corrected fluorescence of widefield fluorescent images for each condition. Significance was calculated by unpaired *t* test with Welch’s correction applied (**** < 0.0001) (POLR2A: N = 142; 6hrActD: N=20; 1hrActD: N=28; DMSO: N= 204). (**C-D**) Representative images from SMLM of EU-labeled RNA for each condition. (**E-G**) Quantification of SMLM distributions for each condition generated using DBscan: fluorophore blink density, cluster density, and cluster size (POLR2A: N = 18; DMSO: N= 28). Significance was calculated by unpaired *t* test with Welch’s correction applied. P-values are listed. (**H-I**) MA plot for POLR2A degraded cells and ActD treated cells showing log2 expression as a function of log fold change using a LFC cutoff of (abs > 1) and signficance level of Padj < 0.05. X axis = Log2 Mean Expression (averaged over all samples). (**J**) Scatterplot of log fold chance values for each gene in one condition compared to log fold change values for the same gene in another condition. Heatmap color scale corresponds to gene density. Marginal distribution shows DEGs with same cutoffs as MA plots. For **B, E– G**: Significance was calculated by unpaired *t* test with Welch’s correction applied (**** < 0.0001).

As anticipated, Pol-II depletion suppressed gene transcription, knocking down many highly expressed genes. However, consistent with our nascent RNA imaging results, depletion of POLR2A triggered upregulation of 5887 genes while only 1959 genes were downregulated. Many of the genes that were upregulated in POLR2A-degraded cells had low basal expression and were not significant initially, undergoing heterogenous increases in expression (**Fig. 2i**). These findings were reproducible and consistent across replicates, indicating that this was not an artifact of library preparation or due to variable mRNA decay rates (**Fig. SI 1i**). Comparing POLR2A to ActD depletion, we were surprised to find that many genes that were downregulated in POLR2A depleted cells were reciprocally upregulated in ActD treated cells and vice versa (**Fig. 2j**). POLR2A degraded replicates had high variance in low expression genes, indicating that its depletion resulted in a significant source of transcriptional heterogeneity without leading exclusively to downregulation. This has also been observed in another recent Pol-II degron experiment^35^. The source of remaining transcription in our system, following Pol-II depletion, may come from active Pol-III or Pol-I or potentially an undegraded subpopulation of Pol-II.

### Transcriptional loops and domain cores are disrupted by Pol-II loss

Given the impact of Pol-II loss on transcriptional patterns genome-wide, we considered that depleting Pol-II may also disrupt the superficial ideal zone of domains where transcriptional reactions are organized, and hence might perturb packing domain architecture directly. To assess if disrupting transcription directly impacted domain architecture, we investigated the effects of POLR2A depletion on chromatin connectivity via Hi-C and *in situ* packing domain structure using a combination of PWS nanoscopy, multiplexed SMLM, and chromatin electron microscopy.

First, we investigated the impact of Pol-II depletion on the formation of forced returns. Previously, we discussed that a subset of these forced returns appear as enhancer-promoter or promoter-promoter loops, visible on Hi-C or Micro-C contact maps. To test this, we generated biological Hi-C replicates for DMSO-treated WT cells (1,094,447,900), POLR2A degraded cells (670,120,527 contacts), and ActD-treated cells (502,274,008 contacts). These replicates were merged to maximize the resolution of each library, producing contact libraries with resolutions below 5000 BP (**SI Fig. 2a**). The individual experiments for each condition were assessed using a stratum-adjusted correlation coefficient and were found reproducible (**SI Fig. 2b)**^36^.

First, we analyzed the impact of transcriptional inhibition on well-established connectivity features such as TADs and compartments. Jiang et. al. previously reported limited impact on TADs and compartments upon degradation of Pol-II^17^. In agreement with previous studies, TADs and A/B compartments do not undergo significant changes following degradation of POLR2A (**Fig. 3e, h**)^17,37,38^ even as ActD-treated cells undergo a change in TADs and compartments (**SI Fig. 3b,d**). We observed little change in genome-wide contact probability at short and long distances (**SI Fig. 3i**).

**Figure 3.**
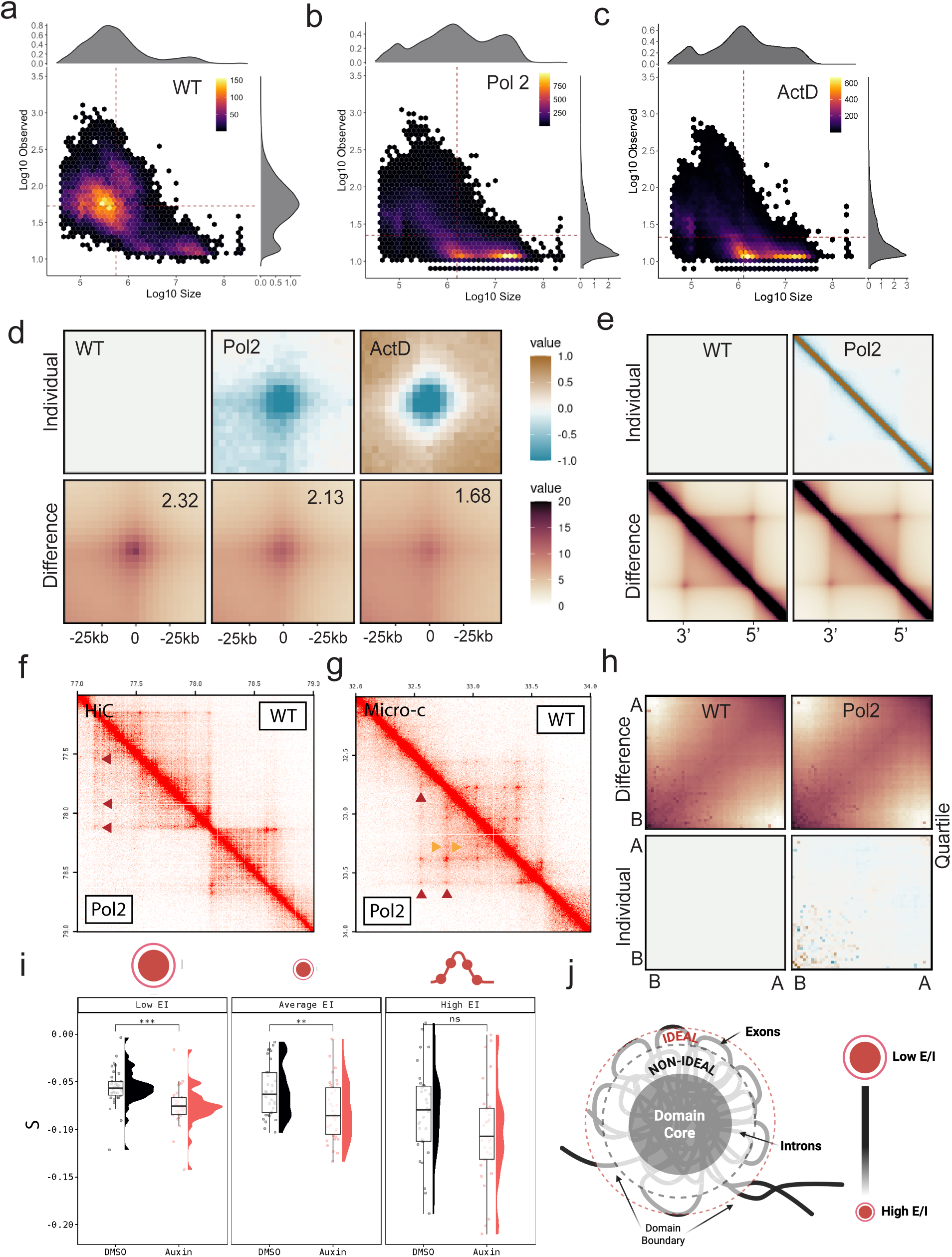
Genome connectivity and *in situ* packing domains are perturbed by Pol-II loss. POLR2A-AID2 degron line was treated with auxin for 6 hours, DMSO for 6 hours, or Actinomycin D treated (5 µg/mL) for 1 hour and Hi-C was generated for each condition. (**A-C**) Scatterplot of log10 loop strength against log10 size for each loop. Loops called using HICCUPS. Heatmap shows loop density (POLR2A Loops: N =56,420; DMSO Loops: N=18,123; ActD Loops: N=39,484). (**D**) Top: difference plot showing change relative to WT. Scale bar: normalized mean loss or gain of contact strength. Bottom: Pileup plots of loop insulation strength for each. Scale bar: normalized mean contact strength. Insulation plots for loop strength were generated in GENOVA^39^. (**E**) TAD insulation plots showing no change to TADs between conditions in POLR2A depleted cells. TADs called using Arrowhead and plotted with GENOVA (**F**) Contact map highlighting loop loss upon POLR2A depletion. Red arrows mark loops that have decreased focal enrichment. (**G**) Contact map from reanalyzed Micro-C data showing distal loop gain and proximal loop loss following POLR2A depletion. Red arrows mark loops that have increased focal enrichment. Yellow arrows mark loops that have decreased focal enrichment. (**H)** Compartment pile-up plot showing no change to compartments (both TAD and compartment pileup plots were generated using GENOVA. See methods for more details). (**I)** Contact scaling for individual genes binned and ranked by E/I score (E/I = Exon Length/(Gene Length – Exon Length) ). Accompanying schematic shows how E/I scales with domain size. Low E/I = large domains, average E/I = small domains, and high E/I = chained aggregates that have are less likely to form domains on their own. **(J)** Schematic demonstrating how the ratio of exons to introns (E/I ratio) within genes organizes domain geometry. Genes with a low E/I ratio are more likely to form domains, wherein heterochromatic introns form sticky cores that exclude euchromatic exons in a transcriptionally poised superficial ideal zone for coherent gene transcription. For **I**: Significance was calculated by unpaired t test with Welch’s correction applied (**** < 0.0001).

We then focused our analysis on loop behavior as a proxy for the impact of transcriptional processes on forced returns. We observed that POLR2A depletion resulted in an increase of many more weak long-range loops (> 1 Mbp), gaining an additional 38,117 loops, even as the loops in wildtype cells become significantly weaker upon POLR2A depletion (**Fig. 3a-b, d**). ActD treatment had a similar effect on loops (**Fig. 3c, d**). Loss of focal enrichment in transcriptional loops following Pol-II depletion was evident in high resolution contact maps (**Fig. 3f**). To validate these results, we analyzed a higher resolution Micro-C dataset generated with the same line from Encode (**Table 5: ENCSR533VUS, ENCSR218URU)** and found that more distal loops were gained while proximal loops experienced decreased enrichment, aligning with other Micro-C studies following transcriptional abrogation where enhancer-promoter and promoter-promoter loops are disrupted (**Fig. 3g**)^20,23^.

One explanation for these observations arises from PD architecture. PDs are comprised of chromatin segments under <2Mbp that are constrained in 3D space due to the combined effects of the excluded volume taken up by chromatin, the spatial constraints of the nucleus, and forced returns. In this schema, if Pol-II generates forced returns to constrain chromatin in space, its depletion will weaken existing transcriptional loops observed on Hi-C and Micro-C. In turn, weakened domain architecture will increase the frequency of unconstrained contacts consistent with the observation of many weak loops (**Fig. 3a-d**). To test this more directly, we took advantage of a recent insight: the surface-to-volume (S/V) assembly of domains may be projected onto the genome from the physical coupling of exons (surface enriched elements) with introns (volume generating elements). This observation suggests that the exon-to-intron (E/I) ratio, especially for large genes (low E/I ratio), can serve as a proxy for the S/V of domains^8^. In this framework, small genes (<50kbp) could either organize outside of domain volumes (as a chain configuration) or share a volume with other small genes. In a constrained volume with well-organized positioning of elements, contact scaling, *S*, is likely to strongly deviate from a polymer relaxed in space (**Fig. 1a-b**). This occurs because, instead of *S* being dominated by the length of the polymer, the contacts between elements are forced together in space. If these constraints are weakened, an observable shift toward a relaxed polymer would occur (trend toward more negative *S*). To test this, we grouped genes by their E/I ratio, binning them into three groups ranging from high E/I to low E/I and measured *S* within genes before and after POLR2A depletion. Consistent with this hypothesis, *S* values within low E/I genes are close to 0 and become more negative upon POL2RA depletion (**Fig. 3i-j**). Notably, the observed transition does not fully return to a relaxed state, suggesting additional constraints on genomic packing independent of Pol- II mediated activity.

A possible mechanism contributing to this behavior could arise from the geometry of domains, where core-forming elements introduce an additional level of physical constraint within packed volumes. For example, although Pol-II may contribute to domain assembly and maintenance, these processes are also co-dependent on orthogonal mechanisms such as heterochromatin-depositing enzymes that stabilize the core of domains^7^. We hypothesized that POLR2A depletion would result in domain core swelling (H3K9me3 spreading out at the nanoscale). To test this hypothesis, we utilized multiplexed SMLM to assay H3K9me3 as a proxy for constitutive heterochromatin cores and POLR2A under different conditions. First, we measured transcriptionally active Pol-II (serine 2 phosphorylated, Pol-II Ps2) in association with packing domain cores, defined by constitutive heterochromatin (H3K9me3). Prior work had demonstrated both that Pol- II Ps2 localized to the transcriptionally active euchromatic periphery of packing domains and that constitutive heterochromatin comprised the dense interior of these domains^7,9,14^. Consistent with these studies, the majority of actively transcribing Pol-II molecules were associated with heterochromatin cores (65%) (**Fig. 4a-b**). Next, we performed SMLM imaging of H3K9me3 alone in POLR2A-depleted vs WT cells.

**Figure 4.**
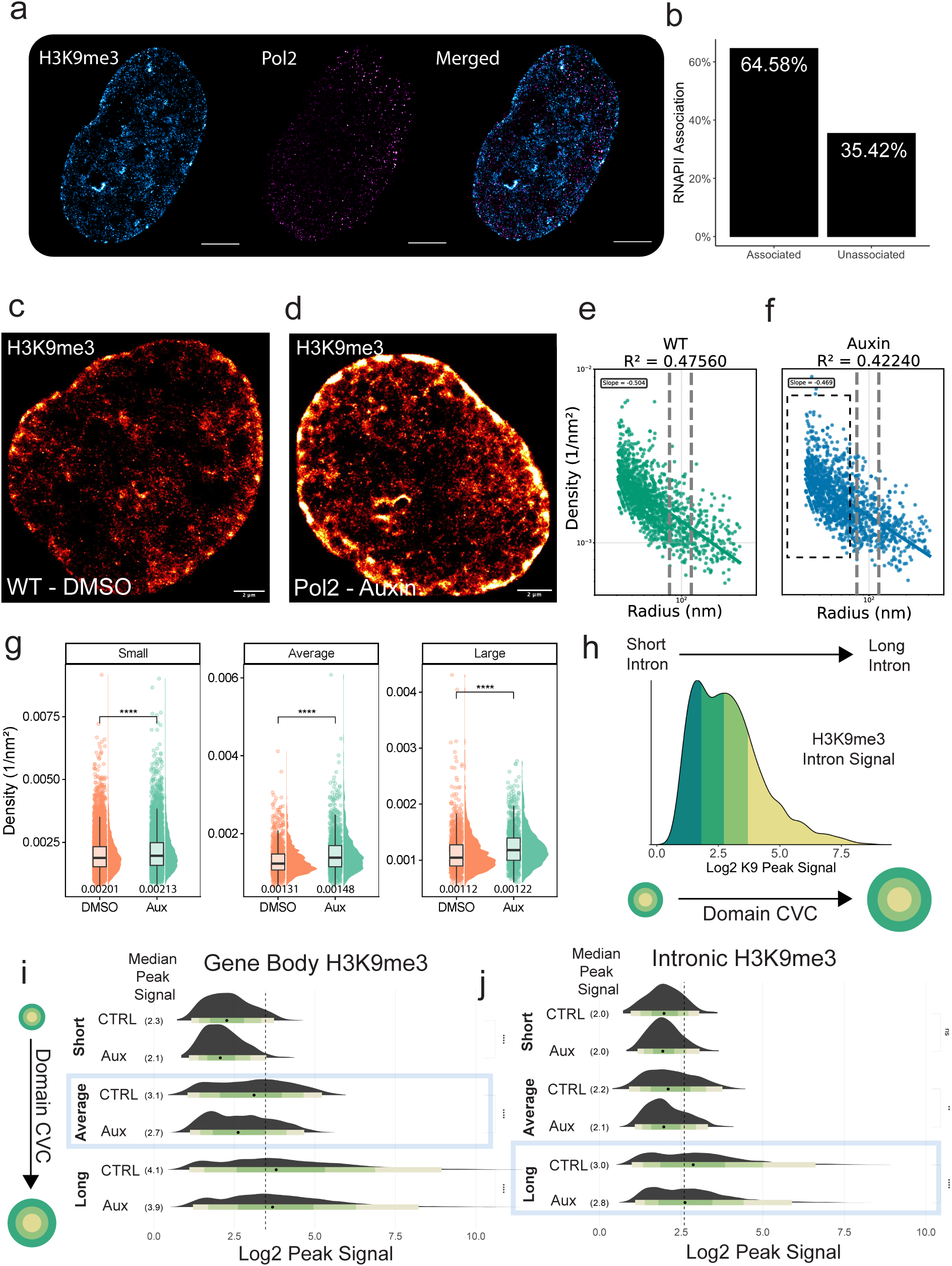
Loss of Pol-II triggers packing domain core degradation and heterochromatin swelling. SMLM data was generated for POLR2A-AID2 line treated with auxin for 6 hours or DMSO for 6 hours (**A**) Representative images of SMLM for labelled H3K9me3 (blue) and labelled active POLR2A (red). (**B**) Association analysis of POLR2A with H3K9me3 clusters showing that POLR2A associates with dense packing domain cores. (**C-D**) Representative images of H3K9me3-labeled SMLM. Scale bar = 2 µm. (**E-F**) Quantification of H3K9me3-conjugated SMLM distributions for each condition: DBscan cluster density and fluorophore blink density, respectively. Data was compiled from two independent biological replicates (DMSO: N = 50; POLR2A: N= 53). (**G**) H3K9me3- conjugated SMLM clusters organized by cluster size and density. In each size category, cluster density increases for auxin treated cells. (**H**) Schematic showing H3K9me3 ChIP-seq peak signal as a function of domain CVC and intron length. Longer introns have more H3K9me3 and contribute to larger domains with a higher CVC. (**I**) Publicly available H3K9me3 ChIP-seq data for a POLR2A-AID2 line treated with auxin for 6 hours or DMSO for 6 hours was analyzed by gene length (**J**) and then by intron length. For **G, I-J,** Significance was calculated by unpaired *t* test with Welch’s correction applied (**** < 0.0001).

On SMLM, the depletion of POLR2A resulted in nuclear-wide reorganization of domain cores. Globally, we observed prominent swelling of heterochromatin at the nuclear periphery (**Fig. 4c-d**). Inhibiting transcription of POLR2A also resulted in a proliferation of larger cores with a higher density of labelled H3K9me3, suggesting that efficient packing at the domain boundaries was breaking down, facilitating antibody penetration to previously sterically excluded regions of the domain (**Fig. 4e-g**) (**SI Fig. 4i**). To verify if this process reflects a disruption of domain geometry, we returned to the S/V packing properties described above to guide analysis of ChIP-Seq data. Whereas ChIP-seq relies on fragmented chromatin for labelling^8^, SMLM may be more sensitive to the sterics of antibody penetration to deeper layers of previously unlabeled chromatin. Therefore, we hypothesized that a partial loss of H3K9me3 within intronic segments on ChIP-seq would be consistent with an ensemble impact on domain packing. We analyzed data from ENCODE for H3K9me3 ChIP-seq with and without POL2RA depletion (**Table 5: ENCSR906ZFR, ENCSR649DZS**). We hypothesized that intron segments forming domain cores are more likely to be enriched in H3K9me3 (**Fig. 4h**)^7,8^. Consistent with our hypothesis, we found that H3K9me3 decreased on introns following Pol-II depletion (**Fig. 4i-j**). Collectively, these results indicate POL2RA is a crucial, but not exclusive, element for domain stabilization. In the absence of forced returns created by Pol-II transcription, domain cores become less compact.

### Gene transcription forms and maintains *in situ* packing domains

To directly interrogate the consequence of disrupting transcription on the physical structure of packing domains, we performed live-cell PWS imaging and ChromSTEM on cells with and without POLR2A depletion. On live-cell PWS, POLR2A depletion resulted in modestly increasing *D_nucleus_* and FMM. The change in FMM can be understood as an increase in the size of nucleosome clutches moving coherently that indicates packing domain deterioration into smaller moving polymeric clutches, while the increase in nuclear *D (D_nucleus_*) suggests more chromatin within PDs (i.e. swelling) (**Fig. 5a-c**). In turn, we observed an acute decrease in the rate of movement of chromatin clutches in POLR2A- degraded cells (**Fig. 5b**), indicating a clear relationship between domain structure and Pol-II activity.

**Figure 5.**
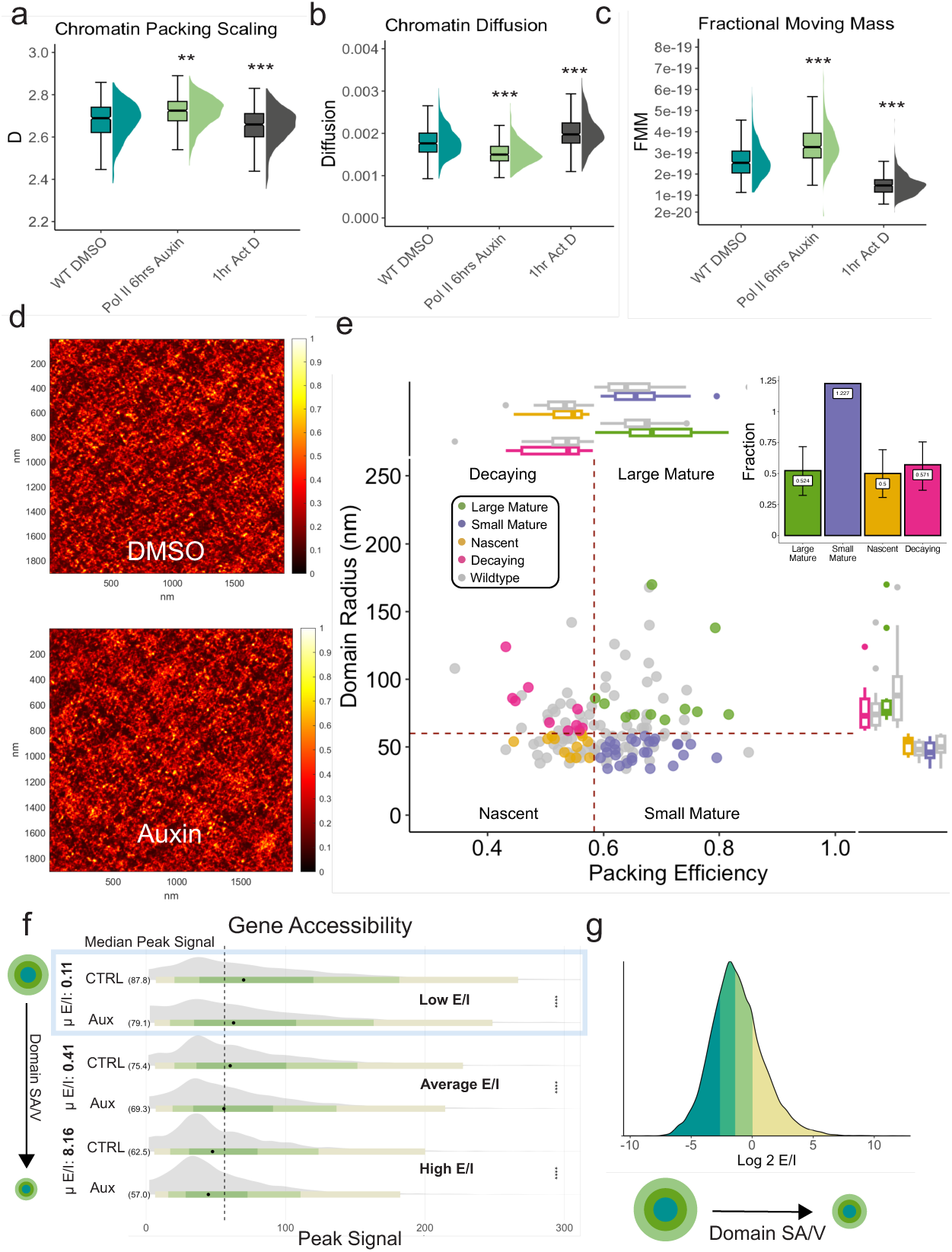
Loss of Pol-II triggers large packing domain degradation and proliferation of small domains. POLR2A-AID2 line was treated with auxin for 6 hours or DMSO for 6 hours and assessed using ChromSTEM. (**A-C**) PWS following treatment: average nuclear packing scaling, diffusion, and fractional moving mass of packing domain nuclear average, respectively. Data was compiled from three independent biological replicates. (DMSO: N = 539; POLR2A: N= 570; ActD: N=475). (**D**) ChromSTEM tomograms showing intact packing domains in untreated cells or degraded packing domain in cells with POLR2A loss. XY axes are in nm. (**E**) Analysis of packing domains by size and packing efficiency to analyze domain properties. Nascent domains (low efficiency, small size), large and small mature domains (high packing efficiency), and decaying domains (low efficiency, large size) represented by color. Lines represent median packing efficiency and median radius in control cells. Each dot represents a single domain. Bar chart in right corner shows number gained or lost in each category (POLR2A: N = 61; DMSO: N=86). (**F**) Publicly available ATAC-seq data for a POLR2A- AID2 line treated with auxin for 6 hours or DMSO for 6 hours was analyzed by Exon/Intron ratio (E/I) for each gene. Exon/Intron ratio corresponds to the surface area to volume ratio (SA/V) of domains (**G**) Schematic showing Log2 E/I as a function of domain SA/V. Low E/I values have large domains and vice versa. For **A-C, F**, Significance was calculated by unpaired *t* test with Welch’s correction applied (**** < 0.0001). For **E**, confidence intervals on bar plot were calculated using a Wilson Score Test.

To measure individual domain structure, we next used ChromSTEM imaging **(Fig. 5d-e)** (**Fig. SI 4a-f**). We categorized domains by their packing efficiency (how well chromatin is packed within the generated volume domains), *⍺*, and domain radius, R. Upon POLR2A depletion, we observed that 48% of large mature domains (>50 nm, well-packed dense domains), 48% of nascent domains (small domains < 50 nm with low density), and 45% of decaying domains (large domains >50 nm with low density) were lost, while small mature domains grew by 23% (**Fig. 5e**)^21^. We next measured the radial density of domains within each respective group and found that the remaining nascent and small mature domains following Pol-II depletion were more compact than small domains in DMSO-treated cells (**Fig. SI 4g**). We then analyzed domain density vs. domain radius and found that mid-density medium (80-120 nm) and large domains (>120 nm) disappeared, but smaller domains (< 80 nm) both proliferated in number and shifted towards an increased density state (**Fig. SI 4h**). These findings were consistent with transcription being involved in domain formation (nascent domain loss) and the stabilization of domain boundaries (loss of large mature and decaying domains). This is in agreement with the loss of heterochromatin observed on ChIP-seq, primarily occurring in long introns (**Fig. 4g-h**).

To measure how this physical transformation translates to the properties of genes, we next utilized publicly available ATAC-seq data (**Table 5: ENCSR182JAI, ENCSR445VYQ)** in the same cell line. We again used E/I to organize genes based on the possible geometry that would satisfy each gene packing into PD volumes. We observed that depletion of POLR2A was associated with a decrease in accessibility per gene (**Fig. 5f-g).** All genes were observed to have a decrease in accessibility but it was particularly prominent for those genes with a low E/I ratio—despite a shift toward smaller and denser domains on chromSTEM—suggests that accessibility scales with PD surface area rather than domain volume. Considering the impact in POLR2A on domain cores (SMLM), PWS imaging, Hi-C, and ChIP-Seq analysis, these findings suggest a crucial role for Pol-II in the formation and maintenance of domain interfaces.

### Active transcription and domain architecture are interdependent

After establishing that transcription and physical structure appear interdependent, we next investigated the functional consequences of disrupting PD architecture. Central to our model is the hypothesis that transcriptional activity generates forced returns— dynamic rearrangement of the chromatin polymer in space—to contribute to domain structure. While ensemble loop analyses from Hi-C and Micro-C provide indirect support for this model, for conclusive support we utilized RNA immunoprecipitation sequencing (RIP-seq), targeting actively elongating polymerase (Pol-II Ps2) to assess whether active transcription is non-randomly distributed along gene bodies. We found that Pol-II Ps2 peaks were enriched over input throughout gene bodies (**SI Fig. 5a**), preferentially occurring at the TES and TSS (**Fig. 6a**). This enrichment pattern is consistent with recent *in vitro* studies demonstrating that transcription-generated supercoiling can drive localized chromatin reorganization, promoting nucleosome compaction behind the transcription bubble and decompaction ahead ^7,8,40,41^. We next mapped Pol-II Ps2 transcripts to introns and exons within gene bodies. Using the method described in Lee et al^42^, the intronic signal was estimated by the difference between the reads mapping to the entire gene body compared to the reads mapping to the annotated exons (**Fig. 6b**). Notably, transcript signal mapped to introns in both input and POLR2A samples decreased toward the intron midpoint, inversely mirroring the coverage profile observed across exons. This pattern aligns with non-uniform polymerase activity along gene architecture and suggests that domain structure may optimize reactions to favor coherent transcription of exons.

**Figure 6.**
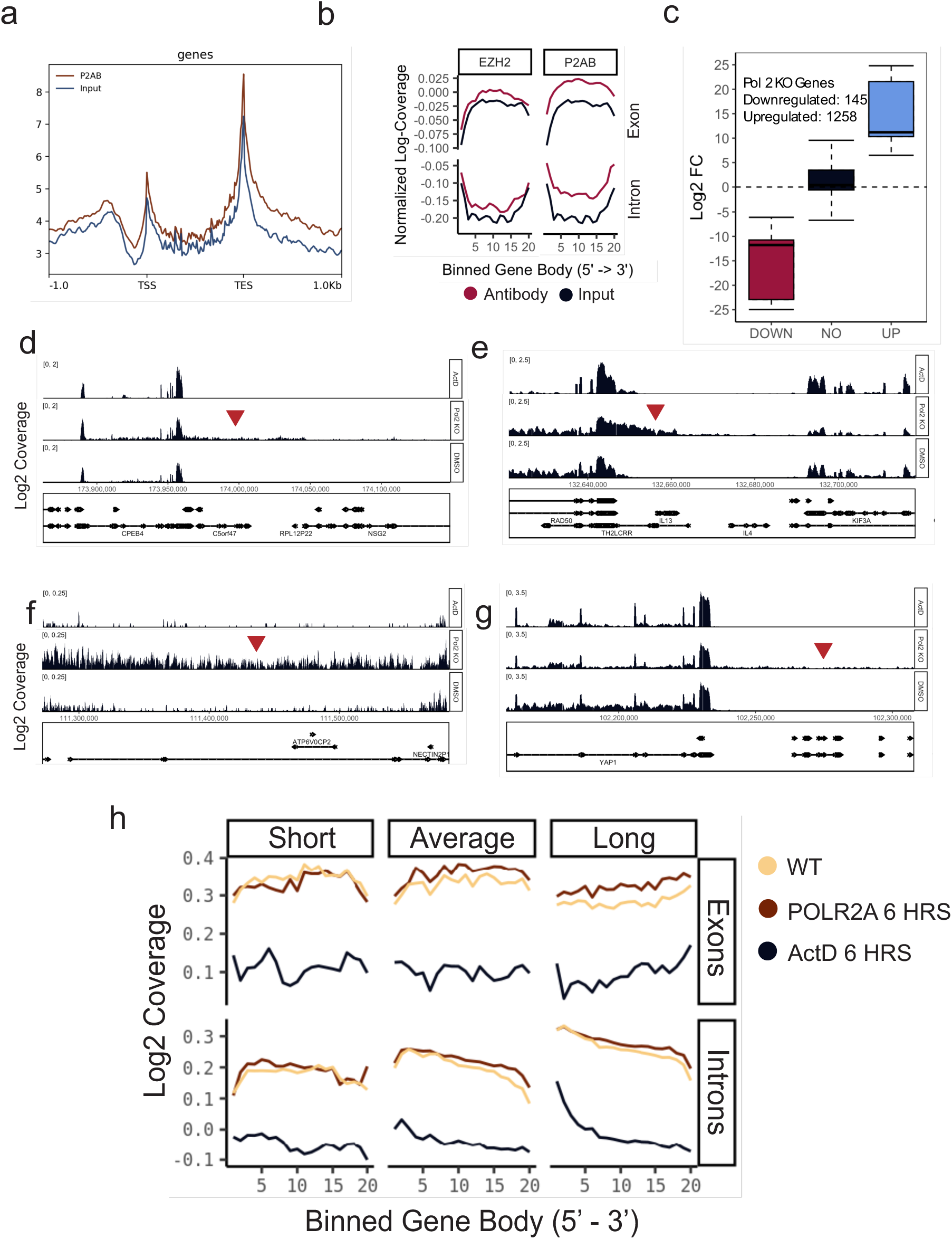
Loss of Pol-II upregulates intronic regions within gene bodies and triggers 5’ readthrough at gene transcription end sites. Analysis of intronic and intergenic expression in RNA-seq generated from POLR2A-AID2 degron line treated with auxin or DMSO for 8 hours or ActD (5 µg/mL) for 6 hours. (**A**) RIP-seq signal on gene bodies for POLR2A immunoprecipitated transcripts in HCT116 cells showing that Pol-II is enriched on the TSS and TES of genes. Region: ± 1kb. (**B**) Intron and exon signal from transcripts immunoprecipitated with an EZH2 (control) or POLR2A antibody in WT HCT116 cells was averaged across 20 bins for all gene bodies. Coverage was normalized, log2 transformed, and plotted. Pol-II RIP in shows gain of signal on both introns and exons. (**C**) Boxplot showing distribution of nascent (EU-seq) differentially expressed genes in POLR2A degraded cells (abs > 0.58) (**D-G**) Representative genomic loci where intron upregulation or 5’ transcriptional readthrough was observed. Red arrows denote 5’ readthrough regions. Orange arrows denote intron upregulation at loci. Coverage was RPGC normalized and Log2 transformed for ease of visualization. (**H**) For nascent transcripts, intron and exon signal was averaged across 20 bins for all gene bodies. Coverage was normalized, log2 transformed, and plotted for each condition by transcript length. ActD shows loss of nascent signal in exons and introns. Pol-II depleted cells show little loss of nascent signal overall, while intron signal decreases with transcript length.

To investigate the functional consequences of Pol-II depletion, we analyzed bases per million (BPM) normalized coverage tracks derived from binary alignment map files for each condition. Coverage scores were log2-transformed (offset by 1 to account for 0- values). We examined genes differentially expressed following POLR2A degradation to explore the aberrant transcriptional behaviors that underlie previous observations (**Fig. 2**). Notably, POLR2A-depleted cells exhibited increased RNA-seq coverage near exon– intron boundaries and elevated intronic signal across gene bodies. In addition, several loci showed extensive 5′ downstream intergenic read coverage. The loss of POLR2A resulted in a mixed population of upregulated and downregulated introns and exons within the same gene, and while the bulk of introns were downregulated, POLR2A loss gave rise to upregulation in many intronic regions (**Fig. 6d–g**). (**SI Fig. 5b**).

Because bulk RNA-Seq reflects steady-state RNA levels, we generated nascent RNA sequencing (EU-seq) to determine the effect of perturbing domain structure on active gene transcription dynamics. In POLR2A degraded cells, most genes were unchanged. However, consistent with the Total RNA-seq results, 1,258 genes were modestly upregulated while 145 genes were downregulated (**Fig. 6c**). We next analyzed transcriptional behavior on introns and exons. We observed that exon transcription was upregulated (N=822) in POLR2A degraded cells compared to controls. Likewise, a small group of introns were robustly upregulated (N=234) (**SI Fig. 5d-e**). In comparison, ActD transcriptional inhibition resulted in suppression of exonic and intronic synthesis (**SI Fig. 5c**). As with our RIP-seq results, nascent transcriptomic RNA is stable from the 5’ to 3’ end of genes on exons but gradually decreases as intron length increases, reflecting domain geometry that optimizes exon transcription and splicing in the ideal zone of domains.

To verify that the 5’ TES readthrough and intron upregulation we observed was reproducible in other data, we analyzed the coverage profiles of the same loci in PRO-seq data for POLR2A depleted HCT-116 cells deposited on ENCODE (**Table 5: ENCSR945WVF**). Consistent with our results, we observed the same readthrough phenotype in this dataset (**SI Fig. 5f-i**). Of these loci, a particularly striking example was the read-through events occurring between RAD50 and IL-13 on Chromosome 5 (**Fig. 6e**) (**SI Fig. 5h**). These genes separately span crucial DNA repair mechanisms (RAD50) and tissue response to inflammatory signals (IL-13 and IL-4)^43–45^. Given the role of IL-13 and IL-4 in inflammatory and atopic disorders, such as asthma, atopic dermatitis, allergic rhinitis, eosinophilic esophagitis, and ulcerative colitis, these results suggest further work is necessary to understand the polymerase-mediated regulatory mechanisms governing chromatin structure between neighboring genes in human disease^7–9^.

### Pol-II degradation does not trigger DNA damage response

We considered that these readthrough events could be the result of pleiotropic effects from destabilization of transcription producing a cellular stress response including downregulation of core splicing machinery or dysregulation of genes in the unfolded protein response (UPR) pathway. To evaluate this alternative, we first quantified TPM-normalized counts in all conditions for core splicing machinery, unfolded protein response, and apoptosis genes. For these categories, 6 hours of Act D treatment was the only condition that saw a substantial drop in total mRNA for essential genes (**SI Fig. 6a**).

Our results indicate that the mRNA for core cellular function genes involved in splicing, UPR, and apoptosis is still stable after 8 hours of polymerase degradation and does not account for the observed readthrough phenotype.

To confirm that Pol-II degradation did not trigger a secondary effect on essential cell homeostasis proteins post-translationally, we generated western blots for p53 (apoptosis), spliced XBP1 (UPR pathway), gamma H2AX (DNA damage response), and Lamin B1 (cell integrity). For each of these, protein levels were unchanged after 6 hours of Pol 2 degradation, affirming that resultant non-coding transcriptional upregulation is not a byproduct of essential pathway destabilization (**SI Fig. 6b-e**).

## Discussion and Conclusion

Many questions persist on the interdependent relationship between supranucleosomal 3D chromatin structure and transcription. Conflicting reports from previous studies depend heavily on the mode of investigation. For example, in older chromosome capture studies, degrading Cohesin or CTCF did not disrupt transcription, while depletion of all three polymerases had little effect on large-scale features such as TADs and compartments, suggesting that topological features and transcription are independent^17–19^. More recent studies using high resolution chromatin capture modalities and nascent RNA-sequencing have demonstrated that loss of TADs does lead to transcriptional and topological dysregulation, but perturbations were limited by the use of pharmacological inhibition and only disrupted fine scale features^20,38^. Large- scale features such as TAD boundaries do exert a limited influence over transcriptional output by insulating-enhancer promoter interactions, however, this behavior is largely probabilistic and non-linear^46^. Pol-II depletion has little impact on TADs and compartments in interphase cells, but Pol-II is necessary to reestablish contact features following exit from mitosis and in embryogenesis^17,37^. Similarly, A and B compartments, long thought to insulate transcriptionally active genes from inactive genes, have both been found to harbor actively transcribing genes using RNA-DNA contact mapping, further muddying the relationship between topological features and gene function^15^. In contrast to topological studies using conformation capture methods, super-resolution STORM imaging found Pol-II distributed along the periphery of packing domains throughout the nucleus^14^, alongside euchromatic post-translational modifications on the outside of heterochromatic cores^9^. And in super-resolution imaging of chromatin, pharmacological disruption of transcription produces a profound impact on 3D chromatin structure^10,11,13^.

To bridge the gap between modes of investigation, we took advantage of an integrative approach in this study previously described by Li et al; we paired *in situ* structural measurements (PWS nanoscopy, multiplexed SMLM, and ChromSTEM) with gene expression analysis, RIP-seq, and Hi-C to resolve the role of transcription in interphase packing domain structure and subsequent function. We show that forced returns generated by RNA polymerase II are required for forming and maintaining PDs as well as for coherent mRNA transcription. Consequently, disrupting Pol-II mediated loops by degrading POLR2A results in dissonant transcriptional signatures where exonic and intronic transcription decouple. While still the primary effect, this does not lead to exclusive gene downregulation, as anticipated. Instead, disrupting Pol-II mediated loops dysregulates expression of exons and introns characterized by frequent readthrough events at gene TES and exon boundaries (**Fig. 6d-g**).

The functional dysregulation described in this study was accompanied by a variety of organizational changes to domains: POLR2A loss precipitated an increase of weak long-range loops and a loss of short range loops (**Fig. 3a-d**). Upon further analysis, we discovered that POLR2A-associated transcripts were specifically enriched on the TSS and TES of genes, supporting a mechanism where Pol-II binding and transcription on either end of gene bodies maintains the physical packing of the domain through torsional force on the chromatin heteropolymer (**Fig. 6a**). In the surrounding organizational context, POLR2A depletion also triggered heterochromatin domain core expansion (H3K9me3) while lowering the likelihood that chromatin would be organized into coherent domains (**Fig. 4c-j**) (**Fig. 5a-c).** Unexpectedly, ground truth measurement of chromatin organization by ChromSTEM tomography revealed that POLR2A loss abruptly triggered large and nascent domain degradation while small mature domains proliferated, consistent with PWS quantification (**Fig. 5d-e**).

While it is possible that an undegraded subpopulation of Pol-II is still active, global transcriptional upregulation on non-coding segments observed after POLR2A depletion raises the possibility that Pol-I or Pol-III are transcribing in Pol-II’s absence. Some recent studies have challenged the paradigm that each polymerase is spatially and functionally restricted. For instance, Pol-II transcription of non-coding RNAs within the 45s rDNA repeat regulate Pol-I rDNA transcription, indicating that the nucleolus is under a co- regulatory framework^47^. While focused on the co-regulatory role of Pol-II and RNA polymerase III (Pol-III), one study found that degrading Pol-I led to differential binding of Pol-II across hundreds of sites^48^. Using RNA-DNA contact mapping, genes actively transcribed by Pol-II were found enriched in contacts to nucleolar organizing hubs^15^. Pol-III consensus binding profiles generated from multiple ChIP datasets predicted widespread Pol-III binding at mRNA promoters^49^. And several studies have found Pol-III transcribing canonical Pol-II genes^48–50^.This opens new lines of inquiry into the mechanisms embedded in packing domain structure that regulate transcriptional coherence and conversely, situations where transcriptional heterogeneity can arise, such as in cancer.

Together, our findings point towards a novel model: Pol-II functions to constrain chromatin in 3D space through generating forced returns in the process of gene transcription. This activity coherently packs the non-coding intronic segments of gene bodies into packing domains, forming nascent domains that evolve through ongoing transcription into packed mature domains. As these domains mature, noncoding elements such as introns function as crucial space-filling scaffolds that form the cores of packing domains; domain cores are then maintained by Pol-II to optimize the physical and molecular conditions for transcription on coding elements near the surface^7,11^. This in turn optimizes the preferred splicing of gene products. In this model, when no longer constrained by Pol-II transcription (Pol-II loss), chromatin becomes unpacked and other polymerases have access to freely transcribe through exon boundaries across unpacked introns and 5’ gene boundaries. The process of unrestricted transcription in turn decreases focal enrichment between promoter-promoter and enhancer-promoter contacts locally, decompacting packing domains and heterochromatin cores in the process (**Fig. 7**).

**Figure 7.**
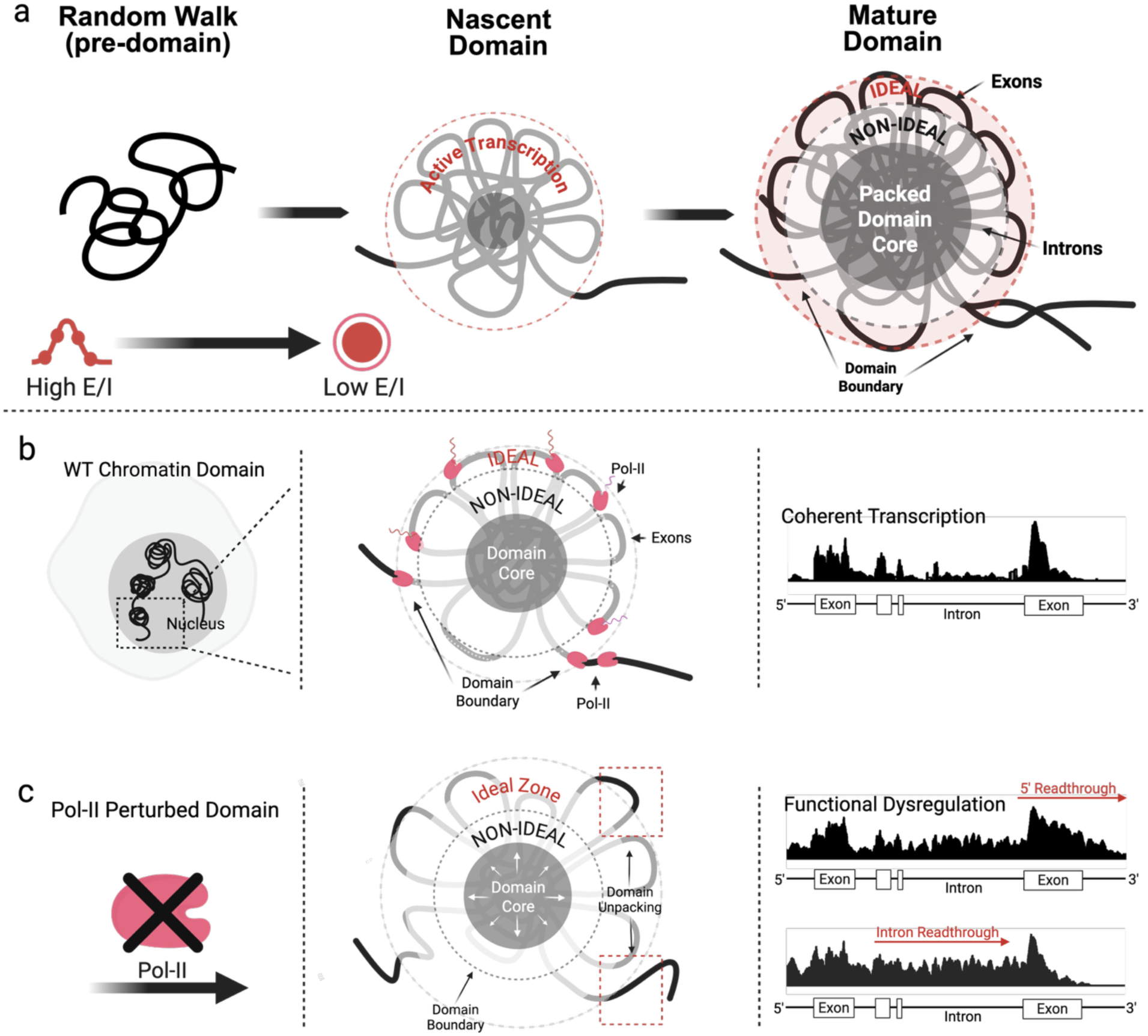
Packing domains are composed of coding and non-coding elements. 98% of the genome is non-coding. Thus, packing domain cores are composed of non-coding intron-rich DNA, held together by “sticky” heterochromatin interactions, where introns are buried within packing domains and sterically excluded from transcriptional proteins while exons are exposed to transcriptional reactions. *Ideal* demarcates zone where transcriptional reactions are favored, near the surface of packing domains, while *non-ideal* demarcates region of domain where transcriptional reactions are not favored. (**A**) Schematic demonstrating the packing domain life cyle: Pol-II forms nascent domains from chromatin engaged in a random walk through the process of transcriptionally driven supercoiling. Nascent domains then mature as chromatin is packed into dense cores. The ratio of exons to introns (E/I ratio) within genes organizes domain geometry. Genes with a low E/I ratio are more likely to form domains, wherein heterochromatic introns form sticky cores that exlude euchromatic exons. (**B**) Model illustrating how transcriptionally driven Pol-II loops constrain packing domain organization and the functional consequence of Pol-II loss. In this model, Pol-II is bound outside of domains and gene boundaries in addition to transcribing along the exons of gene bodies^7–9,51,53,54^. Surface to volume ratio is optimized to favor transcriptional reactions on exons along the ideal zone of domains^8^ (**C**) Packing domain dysregulation upon Pol-II loss leads to heterochromatin core expansion and exposure of intronic sequences to ideal zone, resulting in increased exon readthrough and gain of 5’ gene readthrough into intergenic regions by other polymerases. This also causes domain S/V to increase and domain density to decrease.

With this in mind, new questions arise on the structure-function relationship between conformationally defined packing domains and transcription: 1) How do Pol-II molecules coordinate activity to pack domains into coherent functional units? 2) Do other polymerases act as a short-term failsafe in the absence of Pol-II function? 3) What role do Pol-I and Pol-III play in domain organization, if any? (4) Does packing of intronic and intergenic sequences into domains demarcate genes with distinct functions (RAD50 and IL-13) within domains to coordinate stress responses? For example, the readthrough of RAD50 (associated with DNA damage response) appears to provide a physical basis to couple damage repair to immune recruitment by the expression of IL-13 (a key cytokine involved in allergic and autoimmune disorders). And (5) transcriptional dysregulation is a common feature of multiple cancers—is dysregulation mechanistically associated with cancer transcriptional plasticity and heterogeneity? Supportive prior work suggests that packing domain organization acts as a regulator of an integrated stress response for cells to explore their available genomic information by controlling transcriptional heterogeneity and phenotypic plasticity^51,52^. In line with this, future studies focused on identifying the structural principles governing transcriptional coherence may help uncover new patterns in human diseases. We hope that future work utilizing inducible degron systems, paired studies to other chromatin modifying enzymes, and clinical studies will shed light on these evolving questions.

### Limitations of this study

This study focuses on Pol-II. It is possible that Pol-I and Pol-III also contribute to the transcriptional dysregulation phenotype we observed in our data. Studies going forward should utilize the auxin inducible degron system to target all polymerases. Additionally, although Actinomycin D approximates the phenotype we expect to see when we disrupt all three polymerases, ActD pleiotropy is a poor approximation of polymerase degradation. More complex methods are needed to completely ablate transcription from all three polymerases in a targeted manner and probe the relationship between transcription and chromatin structure. To this end, future studies should generate double degrons targeting pairs of each polymerase.

## Materials and Methods

### Cell Culture

HCT116-POLR2A-AID2^12^ and HCT116 cells were grown in McCoy’s 5A Modified Medium (Thermo Scientific, #16600-082) supplemented with 10% FBS (Thermo Scientific, #16000-044) and penicillin-streptomycin (100 μg/ml; Thermo Scientific, #15140-122). All cells were cultured under recommended conditions at 37°C and 5% CO2. Lines in this study were maintained between passage 5 and 20. Cells were allowed at least 48 hours to re-adhere and recover from trypsin-induced detachment prior to experiments. All experiments and imaging were performed at a confluence between 40–70%. All cells were tested for mycoplasma contamination (ATCC, #30- 1012K) before starting perturbation experiments.

### Auxin Treatment and Drug Treatments

HCT116-POLR2A-AID2 cells were treated with 1 µM 5-phenyl-indole-3-acetic acid (5- Ph-IAA; MedChemExpress, HY-134653) to degrade endogenous POLR2A. HCT116 “WT” cells were treated with either 5 µg/mL Actinomycin D (MP Biomedicals, #0210465810) or % v/v DMSO (Thermo Scientific, #J66650.AE) equal to the highest DMSO concentration in each experiment. Treatment times varied between 1 hour and 8 hours.

### High Throughput Chromatin Conformation Capture

#### Cell Culture and Sample Preparation

Cultured cells were treated with 1 µM 5-PH-IAA or with DMSO at an equivalent concentration for 6 hours. Two technical replicates (treated in separate dishes on the same day) were harvested for each condition. At 6 hours, cells were harvested at 1.2 × 10^6^ cells per sample and transferred to a 50 mL Falcon tube, where they were centrifuged for 10 minutes at 450 x g at room temperature. Supernatant was removed and cells were resuspended in 20 mL of cold fresh media. At this point, 540 µL of 37% formaldehyde (Sigma, #47608) was added to bring the final concentration to 1% formaldehyde and fix the cells for 10 minutes. Upon fixation, the reaction was quenched with 10 mL of 3 M Tris (pH 7.5) (Sigma, #10708976001) for 15 minutes. Cells were then centrifuged for 10 minutes at 800 x g and 4 ° C. After resuspension at the desired concentration, individual samples were snap frozen in liquid N2.

#### Hi-C Library Generation

To wash the nuclei, samples were thawed and resuspended in 50 µL ice cold PB, followed by the addition of 150 µL ice-cold RNase-free water. 50 µL of Buffer C1 (Qiagen Epitect Hi-C Kit, #59971) was added to each sample and mixed. Samples were then centrifuged at 2500 x g and 4 ° C for 5 minutes. The supernatant was aspirated and the nuclear pellet was resuspended in 500 µL of RNase-free water, before being centrifuged again at 2500 x g and 4 ° C for 5 minutes. Library generation and subsequent steps used proprietary reagents from Qiagen’s Epitect Hi-C Kit. Washed nuclei were digested with according to the Epitect Hi-C protocol using a proprietary enzyme cocktail that cut at the GATC motif. Nuclei were end-labeled with biotin followed by ligation for 2 hours at 16 °C. Ligated chromatin was then de-crosslinked using 20 µL of Proteinase K solution at 56 °C for 30 minutes and then heated to 80 °C for 90 minutes. Ligated, decrosslinked DNA was purified using a Qiagen column kit (Qiagen Epitect Hi-C Kit, #59971). and resuspended in 130 µL EB.

#### Library Fragmentation

Hi-C library samples were fragmented to a median size of between 400 and 600 bp using a Covaris E220 sonicator with a sample size of 130 µL and the settings listed in **Table 1**.

**Table 1.** Covaris Parameters:

|  |  |
| --- | --- |
| Water levels | 12 |
| Peak incident power | 140 W |
| Duty factor | 10% |
| Cycles per burst | 200 |
| Treatment time | 55 seconds |

Samples were purified for fragments between 400 and 500 bp using a Qiagen bead purification size exclusion kit (Qiagen Epitect Hi-C Kit, #59971).

#### Hi-C Sequencing Library Generation

Reagents used in Hi-C library generation were supplied with Qiagen Epitect Hi-C Kit unless otherwise noted. Hi-C samples were streptavidin-purified to enrich for properly ligated contact pairs using streptavidin beads and a magnetic bead rack. Beads were first washed in 100 µL of bead wash buffer, resuspended in 50 µL of bead resuspension buffer, and then mixed with 50 µL of Hi-C sample. The mixture was then incubated at room temperature for 15 minutes in a thermal mixer at 1000 RPM. Enriched bead-bound DNA was then end-repaired, phosphorylated, and poly-A tailed using a combined ER/A-tailing solution. The samples were incubated for 15 minutes at 20 °C followed by incubation at 65 °C for 15 minutes. Beads were then washed once with 100 µL of bead wash buffer, washed again with 95 µL of adapter ligation buffer, and resuspended in adapter ligation buffer in preparation for ligation of Illumina adapter sequences. Each sample was mixed with 5 µL of one of 6 Illumina adapter sequences (specified in the Qiagen protocol appendix) and 2 µL of ultralow input ligase. These samples were then incubated for 45 minutes. Following adapter ligation, each sample was washed three times before adding 400 µL of library amplification mixture to the beads. Samples were then distributed equally across 8 wells of a 96-well PCR plate (Thermo Scientific, AB0600L) and cycled using the parameters list in **Table 2** on an Eppendorf Mastercycler x50s thermocycler.

**Table 2.** Hi-C Amplification Parameters:

| Time | Temperature | Cycle No. |
| --- | --- | --- |
| 2 min | 98 ° C | 1 |
| 20 s | 98 ° C |  |
| 30 s | 60 ° C | 7 |
| 30 s | 72 ° C |  |
| 1 min | 72 ° C | 1 |
| hold | 4 ° C | hold |

Following amplification, PCR reactions for each sample were pooled and cleaned using a Qiagen QIAseq library purification kit. Library quality was assessed using a High Sensitivity DNA Assay on an Agilent Bioanalyzer 2100 and a Qubit dsDNA High Sensitivity Assay (Invitrogen, Q33230) Libraries were quantified using a KAPA ROX Low Master Mix qPCR library quantification kit on a QuantStudio 7 Flex instrument. Two samples per lane were sequenced on an Illumina NovaSeq 6000, generating between 400 and 600 million 150 bp paired end reads per sample at Northwestern University’s NUseq Core Facility.

#### Hi-C Data Processing and Analysis

Samples were aligned to human reference genome GRCh38 (Ensembl) using Borrow- Wheeler Aligner version 0.7.17 and processed using the SLURM version of Juicer version 1.6^55^ (https://github.com/aidenlab/juicer) on the Quest HPC provided by Northwestern University. Processing in the Juicer pipeline included removal of duplicates, exclusion of improperly ligated fragments, and mapping of Hi-C contacts with the GATC motif. Statistics generated for each replicate can be found in **SI Fig. 3a**.

Individual replicates were checked for reproducibility using standard heuristics and combined as a mega map using Juicer’s Mega.sh script to increase sample resolution. TADs were identified using Juicer’s Arrowhead algorithm (https://github.com/aidenlab/juicer/wiki/Arrowhead). Loops were identified using Juicer’s HICCUPS algorithm (https://github.com/aidenlab/juicer/wiki/HiCCUPS). Compartment eigenvector analysis and Pearson correlation analysis were generated using Juicer’s Eigenvector and Pearsons scripts, respectively, or using built in functions in GENOVA. Aggregate TAD Analysis and Aggregate Peak Analysis were generated using GENOVA^39^ (https://github.com/robinweide/GENOVA). Contacts were dumped with Hi-C Straw (https://github.com/aidenlab/straw) and used for downstream analysis. Visualization was also done using GENOVA. Differential interaction analysis and cis-trans interaction analysis was done with the MultiHiCompare package in R^56^. Replicate reproducibility was quantified using the HiCRep package in R^36^. All other analyses were done using custom scripts in R.

### Total RNA Library Preparation and Sequencing

Stranded total RNA and small RNA sequencing was conducted in the Northwestern University NUSeq Core Facility. Cultured cells were treated with 1 µM 5-PH-IAA or with DMSO at an equivalent concentration for 8 hours or treated with ActD (5 µg/mL) for 6 hours. Three technical replicates (treated in separate dishes on the same day) were harvested for each condition. At 6 hours, cells were harvested at ∼10 × 10^6^ cells per sample and transferred to 1.7 mL Eppendorf tubes. Samples were centrifuged and media was aspirated, followed by snap freezing in N2. Later, total RNA was extracted with the miRNeasy kit (Qiagen, #217084). Total RNA examples were checked for quality on Agilent Bioanalyzer 2100 and quantity with Qubit fluorometer. For sequencing of large RNA species, the Illumina Stranded Total RNA Library Preparation Kit (Illumina, 20040525) was used to prepare sequencing libraries. The kit procedure was performed without modifications. This procedure includes rRNA depletion, remaining RNA purification and fragmentation, cDNA synthesis, 3’ end adenylation, Illumina adapter ligation, library PCR amplification and validation. For sequencing of small RNAs, the NEXTFLEX Small RNA-Seq Kit v4 (Revvity, NOVA-5132-32) was used to build sequencing libraries. First, 3’ RNA adapter and then 5’ adapter were ligated to microRNAs and other small RNAs in the samples. After ligation, a reverse transcription step was carried out to generate single-stranded cDNA. A PCR step is then conducted to amplify the cDNA and incorporate Illumina adapter sequences including index sequences. The amplified cDNA constructs were then purified with a bead cleanup step prior to sequencing. Illumina NovaSeq X Plus and HiSeq 4000 NGS Systems were used to generate single 50-base reads for each sample.

### RNA-seq Data Analysis

RNA-seq reads were preprocessed with FastQC v.0.12.0 and aligned with STAR v.2.6.0^57^ using the --quantMode TranscriptomeSAM parameter to human reference genome GRCh38 (Ensembl). Aligned samples were converted to binary alignment map (BAM) files, sorted, and indexed using SAMtools v.1.6^58^. Coverage files were generated as BigWigs for the postive and negative strand using DeepTools v.3.1.1^59^ BamCoverage utility and normalized by either CPM or RPGC. Coverage files were later merged for ease of analysis. Alignment statistics were generated using SAMtools Flagstat utility. Aligned and processed BAMs were then quantified to generate a counts table using the rsem-calculate-expression command from RSEM v.1.3.3^60^ or the htseq-count command from HTSeq v.2.0.2^61^ and the Homo_sapiens.GRCh38.111.gtf (Ensembl) genome annotation. Differential gene expression analysis was performed using DESeq2 v.1.44.0 in R. Intron-centric differential expression analysis was performed using the Intron Differences To Exon or INDEX package (https://github.com/Shians/index) and Superintronic package (https://github.com/sa-lee/superintronic) in R^42^. Visualizations of coverage files were generated using the Gviz^62^ package in R.

### EU-seq

Cultured cells were treated with 1 µM 5-PH-IAA or with DMSO at an equivalent concentration for 8 hours or treated with ActD (5 µg/mL) for 6 hours. Following treatment, nascent RNA was labelled, captured, and sequenced following guidance from [Palozola 2021](https://www.sciencedirect.com/unsupported_browser). To selectively isolate EU-labeled nascent RNA, HCT116 cells were treated with media containing 0.5 mM 5-ethynyluridine (EU) for 1 hour (Click-iT nascent RNA Capture Kit cat. no. c10365, Thermo Fisher Scientific). These RNAs were retrieved using Click-iT chemistry to bind biotin azide to the ethylene group of EU-labeled RNA. The EU-labeled nascent RNA was purified using MyOne Streptavidin T1 magnetic beads. Captured EU-RNA attached on streptavidin beads was immediately subjected to on-bead sequencing library generation using the Universal Plus Total RNA-Seq with NuQuant® (Tecan) kit according to the manufacturer’s protocols with modifications. On-bead complementary DNA (cDNA) was synthesized by reverse transcriptase using random hexamer primers. The cDNA fragments were then blunt-ended through an end-repair reaction, followed by dA-tailing. Subsequently, specific double-stranded barcoded adapters were ligated and library amplification for 15 cycles was performed. PCR libraries were cleaned up, measured on an Agilent Bioanalyzer using the DNA1000 assay, pooled at equal concentrations and sequenced on a Novogene Nova Seq X with 50 BP paired end.

### EU-seq Data Analysis

Paired end nascent RNA transcripts were trimmed with TrimGalore v0.6.10 and aligned using Hisat2 v2.1.0. Alignment files were sorted, filtered, and converted to BAM format using Samtools v1.6. Coverage files were generated using Deeptools v3.1.1 and normalized using RPGC or BPM normalization. Finally, aligned filtered reads were counted using separate genome annotation files for intron, exon, gene bodies, and intergenic regions with Htseq v2.0.2. Genome annotations for introns and exons were generated using the methods described by Alkallas & Najafabadi ([Alkallas & Najafabadi 2022](https://link.springer.com/protocol/10.1007/978-1-0716-2409-8_9)). An intergenic genome annotation was generated using a custom script by subtracting the union of exons and introns from gene bodies and dividing the remainder into countable bins of 25kb each. Gene body coverage plots were generated using the Superintronic package described in [Lee et al 2020](https://academic.oup.com/nargab/article/2/3/lqaa073/5910008) by binning each gene body into 20 bins in the 5’->3’ orientation and calculating the average coverage in each bin for short, average, and long genes within introns and on exons. For differential expression analysis, exons and introns with signal in the EU control were subtracted from the analysis.

### RIP-seq

Cultured cells were treated with 1 µM 5-PH-IAA or with DMSO at an equivalent concentration for 5 hours. Following auxin or DMSO treatment, protein-bound RNAs were immunoprecipitated using the EZ- Magna™ Nuclear RIP (Native) kit (Cat. # 17-10521) kit as follows. Cells were spun down and resuspended in ice cold resuspension buffer, homogenized, and lysed. Lysates were treated with DNAse I. Inputs for each sample were withdrawn at this stage. Lysates were then incubated with 5 µg of protein A/G Magnetic Beads bound primary antibody for the protein of interest and incubated on a rotating rack and incubate at 4°C for overnight. Following the immunoprecipitation step, lysates were again treated with DNAse I before going through a wash step. After wash steps, RNA was purified from IPs using Trizol reagent and chloroform per the protocol. Purified RNAs were processed as a library using the same library generation kit that was used for the bulk RNA-sequencing. rDNA was removed for all samples except for one sample from each condition set aside for rDNA quantification. The Illumina HiSeq 4000 NGS System was used to generate single 50-base reads for each sample.

### RIP-seq Data Analysis

Paired end nascent RNA transcripts were trimmed with TrimGalore v0.6.10 and aligned using Hisat2 v2.1.0 to human reference genome GRCh38 (Ensembl). Duplicate reads were marked and removed using Picard v.2.21.4^63^. Alignment files were sorted, filtered, and converted to BAM format using Samtools v1.6. Coverage files were generated using Deeptools v3.1.1 and normalized using RPGC or BPM normalization. BAMs were merged for final consensus peak calling. Peaks were called on BAMs with MACS2 v. 2.2.9.1^64^ using the -q 0.1 --keep-dup all parameters and with Pirhana ([Uren 2013](https://pmc.ncbi.nlm.nih.gov/articles/PMC3509493/)). Pirhana peaks were called using a q value cutoff of 0.1 and a binsize of 200 BP. Finally, aligned filtered reads were counted using separate genome annotation files for intron, exon, gene bodies, and intergenic regions with Htseq v2.0.2 with the Homo_sapiens.GRCh38.111.gtf (Ensembl) genome annotation. Genome annotations for introns and exons were generated using the methods described by Alkallas & Najafabadi ([Alkallas & Najafabadi 2022](https://link.springer.com/protocol/10.1007/978-1-0716-2409-8_9)). An intergenic genome annotation was generated using a custom script by subtracting the union of exons and introns from gene bodies and dividing the remainder into countable bins of 25kb each. Gene body coverage plots were generated using the Superintronic package described in [Lee et al 2020](https://academic.oup.com/nargab/article/2/3/lqaa073/5910008) by binning each gene body into 20 bins in the 5’->3’ orientation and calculating the average coverage in each bin for short, average, and long genes within introns and on exons. Matrices of peak coverage profiles were generated using Deeptools v.3.1.1 computeMatrix and heatmaps were plotted using plotHeatmap. Differential gene expression analysis was performed using DESeq2 v.1.44.0 in R. Input vs coverage plots were generated by taking the overlap of coverage signal and RIP-seq peaks called with MACS2 in the treatment condition. Intron-centric differential expression analysis was performed using the Intron Differences To Exon or INDEX package (https://github.com/Shians/index) and Superintronic package (https://github.com/sa-lee/superintronic) in R^42^. Visualizations of coverage files were generated using the Gviz^62^ package in R.

### Multi-Color and Single Color SMLM Sample Preparation

#### For H3K9me3 Single Label Imaging

1. Cells were plated on No. 1 borosilicate bottom eight-well Lab-Tek Chambered cover glass (Thermo Scientific, #155411) at a seeding density of 1∼2.5k. After 48 hours, the cells underwent fixation for 10 mins at room temperature with a fixation buffer composed of 4% paraformaldehyde (Electron Microscopy Sciences, #15710) in PBS. Samples were then washed in PBS for 5 minutes and quenched with freshly prepared 0.1% sodium borohydride (Sigma, # 213462) in PBS for 7 min. Two more wash steps were performed after quenching.
2. Permeabilization was done with blocking buffer composed of (3% bovine serum albumin (BSA) (Fisher, #BP671-10), 0.5% Triton X-100 (Thermo #A16046-AE) in PBS) for 1 hour and then samples were immediately incubated with rabbit anti-H3K9me3 antibody (Abcam, ab176916) in blocking buffer for 1-2 hours at room temperature on an orbital shaker. Samples were then washed three times with a washing buffer composed of 0.2% BSA and 0.1% Triton X-100 in PBS.
3. Afterwards, samples were incubated with the corresponding goat antibody–dye conjugates, anti-rabbit AF647 (ThermoFisher, A-21245) for 40-60 mins at room temperature with shaking. After incubation, samples were washed two times in PBS for 5 mins with shaking followed by imaging.

#### For H3K9me3 and RNAP polymerase II Dual-Label Imaging

Steps 1,2 and 3 were repeated as described above. Samples were then incubated overnight in a modified version of the blocking buffer, comprised of 10% goat serum (ThermoFisher, 16210064) and 90% the prior composition. After overnight blocking, the samples would then go through the same protocol as in step 3 (primary and secondary antibody incubation) but this time with modified blocking buffer (90% original blocking buffer + 10% goat serum) and washing buffer (99% original washing buffer + 1% goat serum) for the second target. The second target primary antibody was rat anti-RNA Polymerase II antibody (Abcam, ab252855) and the secondary antibody was goat anti-rat AF488 (ThermoFisher, A-150157). As described before, samples were blocked overnight between labels at 4°C. After the primary and secondary incubation, the samples were washed three times with PBS followed by imaging. All antibodies used in this study are listed on **Table 3**.

**Table 3.**
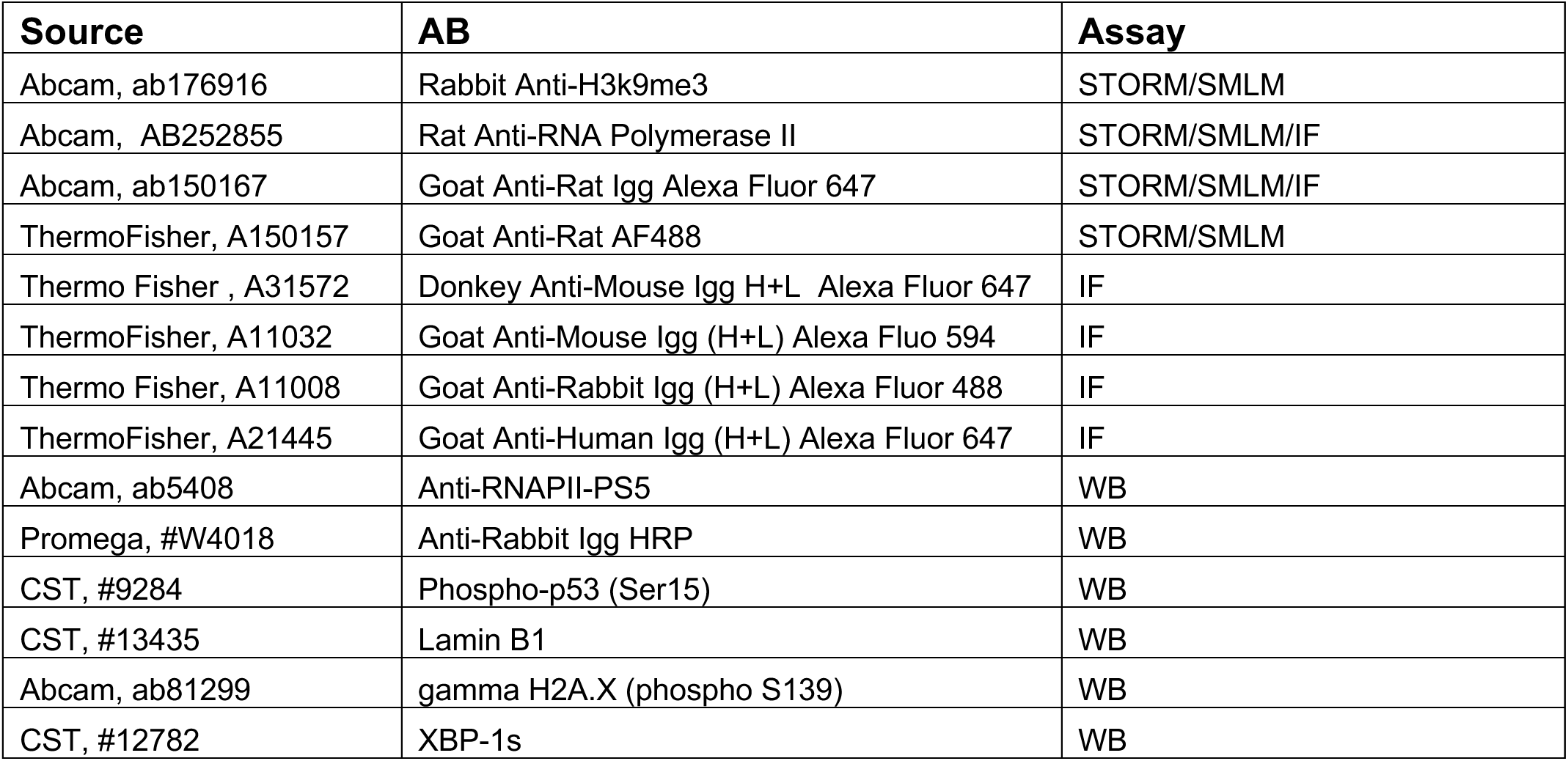
Antibodies:

| Source | AB | Assay |
| --- | --- | --- |
| Abcam, ab176916 | Rabbit Anti-H3K9me3 | STORM/SMLM |
| Abcam, AB252855 | Rat Anti-RNA Polymerase II | STORM/SMLM/IF |
| Abcam, ab150167 | Goat Anti-Rat IgG Alexa Fluor 647 | STORM/SMLM/IF |
| ThermoFisher, A150157 | Goat Anti-Rat AF488 | STORM/SMLM |
| Thermo Fisher, A31572 | Donkey Anti-Mouse IgG H+L Alexa Fluor 647 | IF |
| ThermoFisher, A11032 | Goat Anti-Mouse IgG (H+L) Alexa Fluor 594 | IF |
| Thermo Fisher, A11008 | Goat Anti-Rabbit IgG (H+L) Alexa Fluor 488 | IF |
| ThermoFisher, A21445 | Goat Anti-Human IgG (H+L) Alexa Fluor 647 | IF |
| Abcam, ab5408 | Anti-RNAPII-PS5 | WB |
| Promega, #W4018 | Anti-Rabbit IgG HRP | WB |
| CST, #9284 | Phospho-p53 (Ser15) | WB |
| CST, #13435 | Lamin B1 | WB |
| Abcam, ab81299 | gamma H2A.X (phospho S139) | WB |
| CST, #12782 | XBP-1s | WB |

#### Single Molecule Localization Data Analysis

Acquired data was first processed using the ThunderSTORM ImageJ plugin^65^ to generate the reconstructed images for visualization via the average shifted histogram method, as well as the localization datasets. Each localization dataset was corrected for drift and subsequently filtered such that remaining data had an uncertainty of less than or equal to 40 nm. Localization coordinates (x,y) were then used in a Python point cloud data analysis algorithm which employed the scikit-learn DBSCAN method (min_pts=3, epsilon=50) to cluster the heterochromatic localizations. Cluster size was determined by the area of the Convex Hull fit of the clustered marks and then normalized relative to a circular cluster with radius of 80 nm. POLR2A density was measured by counting the number of POLR2A in dilated contours of the identified cluster periphery (rings that follow the shape of the cluster) and dividing by the ring area. Association was determined by measuring the number of POLR2A that fall within 5 times the area outside of the cluster relative to all POLR2A localizations. The outside cluster condition signifies and POLR2A that is not within any analysis area and thus is not associated with heterochromatic clusters.

### Immunofluorescence Imaging

HCT116-POLR2A-AID2 cells were plated at 10,000 cells per well of an 8-chamber cover glass plate (Cellvis C8-1.5H-N). Following auxin treatment, cells were washed twice with 1x Phosphate Buffered Saline (PBS) (Gibco, #10010031). Cells were fixed with 4% paraformaldehyde (PFA) (Electron Microscopy Sciences, #15710) for 10 minutes at room temperature, followed by washing with PBS 3 times for 5 minutes each. Cells were permeabilized for 15 minutes using 0.2% TritonX-100 (10%) (Sigma-Aldrich, #93443) in 1x PBS, followed by another wash with 1x PBS for 3 times for 5 minutes each. Cells were blocked for 1 hour using 3% BSA (Sigma-Aldrich, #A7906) in PBST (Tween-20 in 1x PBS) (Sigma-Aldrich, #P9416) at room temperature. The following primary antibodies were added overnight at 4°C: Rat POLR2A PS2 antibody (ABCam, AB252855). Cells were washed with 1x PBS 3 times for 5 minutes each. Either of the following secondary antibodies were added for 1 hour at room temperature: Goat Anti-Rat IgG Alexa Fluor 647 (Abcam, ab150167, dilution 1:1000) or Donkey Anti-Mouse IgG H+L Alexa Fluor 647 (Thermo Fisher Scientific, A31572, dilution 1:500). Cells were washed with 1x PBS 3 times for 5 minutes each. Finally, cells were stained with DAPI (Thermo Fisher Scientific, #62248, diluted to 0.5 μg/mL in 1x PBS) for 10 minutes at room temperature. Prior to imaging, cells were washed with 1x PBS twice for 5 minutes each. The cells were imaged using the Nikon Eclipse Ti2 inverted microscope equipped with a Prime 95B Scientific CMOS camera. Images were collected using a 100x/ 1.42 NA oil-immersion objective mounted with a 2.8x magnifier. mClover was excited at 488 nm laser, Alexa Fluor 647 was excited with at 633 nm, and DAPI was excited at 405 nm. Imaging data were acquired by Nikon acquisition software. All antibodies used in this study are listed on **Table 3**.

### EU-labeled nascent RNA staining and imaging

For EU-labeled nascent RNA staining and imaging in vivo, we used the Click-iT RNA Alexa Fluor 647 Imaging Kit (Thermo, C10330) according to the standard immunofluorescence protocol. Cells were plated at 50,000 per well in a 12-well optical plate (Cellvis, P12-1.5H-N) for widefield fluorescence imaging or at 10,000 cells per chamber in an 8-chamber cover glass plate (Cellvis C8-1.5H-N). 48 hours after plating, partially confluent dishes were treated as described above to degrade POLR2A or all three polymerases (5 µg/mL Actinomycin D). 1 hour prior to fixation, 1 mM EU was added to each well. Cells were washed with PBS and fixed with 4% paraformaldehyde for 10 minutes followed by three more washes in 3% bovine serum albumin (BSA) PBS and permeabilization using 0.5% Triton X-100 in PBS for 20 min at room temperature. Each well was then washed twice with 3% BSA in PBS and incubated with Click-iT reaction mix for 30 min. After the Click-iT reaction, cells were washed twice with 3% BSA in PBS. For widefield epifluorescence, all conditions were imaged using the Nikon Eclipse Ti2 inverted microscope equipped with a Prime 95B Scientific CMOS camera. For EU-STORM, the steps were the same but all conditions were imaged using the SMLM parameters described previously.

### Data and Image Analysis

For immunofluorescence and widefield fluorescence experiments, images were analyzed through imaging processing pipelines utilizing standard tools in Fiji v.2.14.0^66^. Corrected Total Cell Fluorescence (CTCF) intensity was quantified by subtracting out background signal and correcting for intensity differences due to area as previously described, CTCF = Integrated Density – (Area of selected cell X Mean fluorescence of background readings)^67^.

### ChromSTEM

#### Electron Microscopy Sample Preparation

The chemical reagents are listed on **Table 4a** and **Table 4b**. Cells were fixed with 2% paraformaldehyde, 2.5% glutaraldehyde (EM-Grade), 2mM CaCl_2_ in 0.1M sodium cacodylate buffer for 30 minutes at room temperature and 30 minutes at 4°C. The cells were then kept in cold temperature, if possible, for further treatments. After fixation, cells were washed 5x for 2 minutes each wash with 0.1M sodium cacodylate buffer and blocked with 10mM glycine, 10mM potassium cyanide, 0.1M sodium cacodylate buffer for 15 minutes.

**Table 4a.** Reagents used in ChromSTEM Staining:

| Reagent | Formula |
| --- | --- |
| Washing solution | Hank's balanced salt solution without calcium and magnesium |
| Fixation solution | 2.5% EM grade glutaraldehyde<br>2% paraformaldehyde<br>2 mM CaCl<sub>2</sub><br>0.1 M sodium cacodylate buffer, pH = 7.4 |
| Blocking solution | 10 mM glycine<br>10 mM potassium cyanide<br>0.1 M sodium cacodylate buffer, pH = 7.4 |
| DNA staining solution | 10 µM DRAQ5<br>0.1% SAPONIN<br>0.1 M sodium cacodylate buffer, pH = 7.4 |
| Bathing solution | 2.5 mM 3,3'- diaminobenzidine tetrahydrochloride (DAB)<br>0.1 M sodium cacodylate buffer, pH = 7.4 |
| Reduced osmium staining solution | 2% osmium tetroxide<br>1.5% potassium ferrocyanide<br>2 mM CaCl<sub>2</sub><br>0.15 M sodium cacodylate buffer, pH = 7.4 |
| Durcupan™ resin mixture 1 | 10 mL Durcupan™ ACM single component A, M, epoxy resin<br>10 mL Durcupan™ ACM single component B, hardener 964<br>0.15 mL Durcupan™ ACM single component D |
| Durcupan™ resin mixture 2 | 10 mL Durcupan™ ACM single component A, M, epoxy resin<br>10 mL Durcupan™ ACM single component B, hardener 964<br>0.2 mL Durcupan™ ACM, single component C, accelerator 960<br>0.15 mL Durcupan™ ACM single component D |
| 1:1 infiltration mixture | 10 mL 100% ethanol<br>10 mL Durcupan™ resin mixture 1 |
| 2:1 infiltration mixture | 5 mL 100% ethanol<br>10 mL Durcupan™ resin mixture 1 |

**Table 4b.**
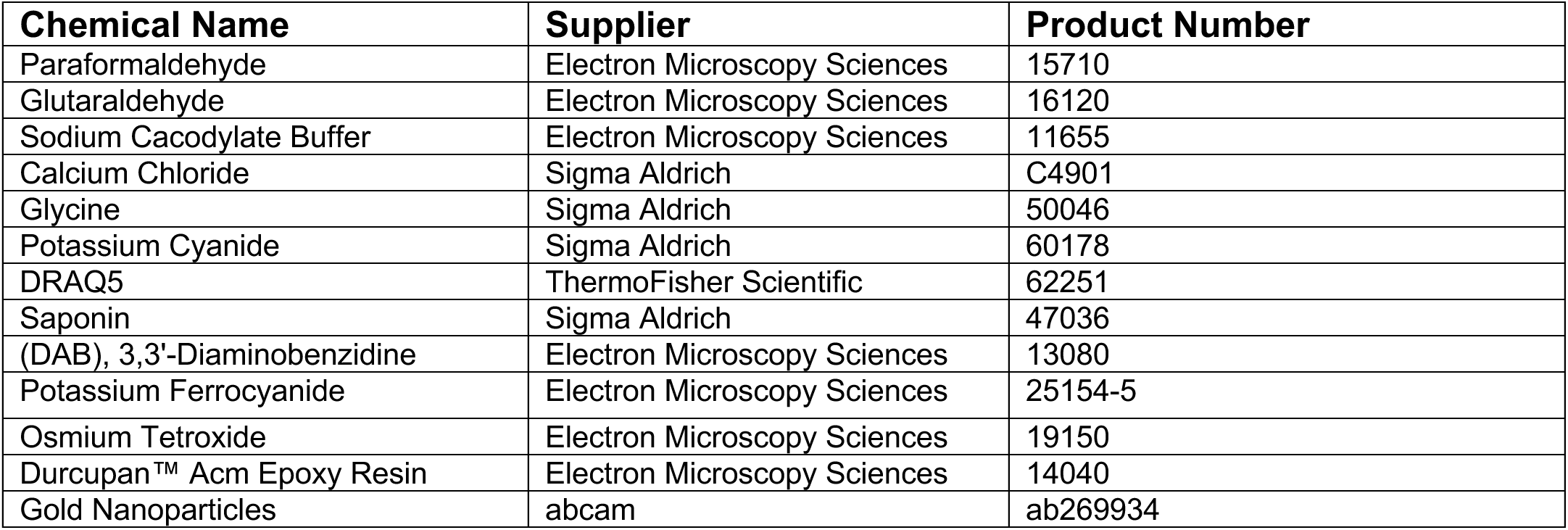
Reagents used in ChromSTEM Staining:

| Chemical Name | Supplier | Product Number |
| --- | --- | --- |
| Paraformaldehyde | Electron Microscopy Sciences | 15710 |
| Glutaraldehyde | Electron Microscopy Sciences | 16120 |
| Sodium Cacodylate Buffer | Electron Microscopy Sciences | 11655 |
| Calcium Chloride | Sigma Aldrich | C4901 |
| Glycine | Sigma Aldrich | 50046 |
| Potassium Cyanide | Sigma Aldrich | 60178 |
| DRAQ5 | ThermoFisher Scientific | 62251 |
| Saponin | Sigma Aldrich | 47036 |
| (DAB), 3,3'-Diaminobenzidine | Electron Microscopy Sciences | 13080 |
| Potassium Ferrocyanide | Electron Microscopy Sciences | 25154-5 |
| Osmium Tetroxide | Electron Microscopy Sciences | 19150 |
| Durcupan™ Acm Epoxy Resin | Electron Microscopy Sciences | 14040 |
| Gold Nanoparticles | abcam | ab269934 |

**Table 5.**
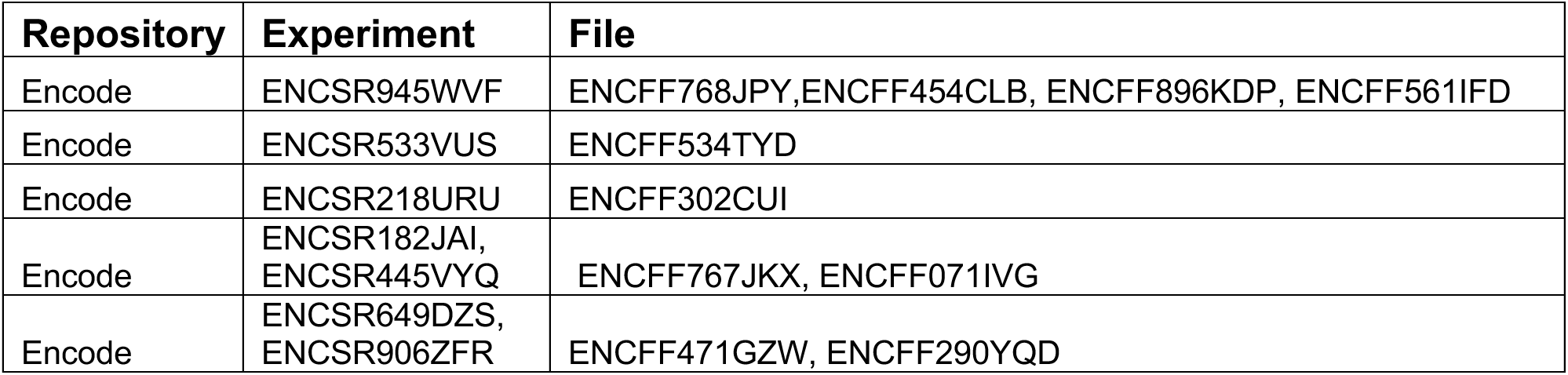
Publicly Available Data Utilized in this Study:

The cells were stained with 10μM DRAQ5, 0.1% saponin, 0.1M sodium cacodylate buffer for 10 minutes, followed by 3x washes for 5 minutes each with the blocking buffer. After that, the cells were photo-oxidized in 2.5mM 3,3’-Diaminobenzidine (EM- Grade) under a 100X oil objective, 15W Xenon lamp and Cy5 filter for 5 minutes. The cells were washed 5x times for 2 minutes each with 0.1M sodium cacodylate buffer and stained with 2% osmium tetroxide, 1.5% potassium ferrocyanide, 2mM CaCl_2_ in 0.15M sodium cacodylate buffer for 30 minutes. Cells were washed 5x times for 2 minutes each with Millipore water afterwards.

Cells were then dehydrated gradually with 30%, 50%, 70%, 85%, 95%, 2 times 100% ethanol. After that, cells were incubated under room temperature with 100% ethanol, followed by embedding with Durcupan resin 1:1 infiltration mixture for 30 minutes, 2:1 infiltration mixture overnight, resin mixture 1 for 1 hour, resin mixture 2 for 3 hours. Lastly, the samples were embedded with resin mixture 2 at 60° C for 48 hours and were collected for ultramicrotomy.

For ultramicrotomy, an ultramicrotome (UC7, Leica) along with a 35° Diatome knife were used to section 120 nm thick resin samples. The sections were collected on a copper slot grid (2 x 0.5mm) with formvar/carbon film. Gold nanoparticles with 10nm diameter were deposited on both surfaces of the grid afterwards as fiducial markers.

#### Image Collection and Tomography Reconstruction

Images were collected with Hitachi HD2300 STEM microscope at 200kV with HAADF imaging mode, at a magnification of 50kX. Two sets of tilt series images were collected by rotating samples from -60° to +60° at a 2° step, with two roughly perpendicular rotation axes.

For post-processing, IMOD was used to align the images. The gold nanoparticles of the collected images were removed with IMOD for another set of data without the influence of extreme values from gold nanoparticles. For each tilt series, Tomopy was used to reconstruct the volume with Penalized maximum likelihood algorithm with weighted linear and quadratic penalties. The two independent reconstructed volumes were then combined in IMOD, with gold nanoparticles as the matching model and repeated on the data without nanoparticles.

After reconstruction, extreme pixel values (1.5 times standard deviation below the mean and 5 times standard deviation above the mean) were capped. The pixel values were then scaled between 0 to 1 for analysis.

### Packing domain Identification and Analysis

Packing domains were identified and analyzed following the approach previously described^6^.

For identification of domains, a Gaussian filter with radius = 5 pixels was used followed by CLAHE contrast enhancement on the 2D projection of the 3D tomogram using ImageJ. Packing domain centers were identified as local maxima with prominence = 1.5 x standard deviation of pixel values.

To quantify domain properties, an 11 x 11 pixels window is applied to each domain and sampled for each domain. Packing scaling analysis is done by measuring the total intensity of the chromatin that radially expands from the center pixel picked and weighted by the intensity of the center pixel. The linear region of the packing scaling behavior is identified by MATLAB ‘ischange’ function. The domain size is measured as the point at which the packing scaling behavior deviates from the linear fits with 5% difference, or when local packing scaling exponent D reaches 3. CVC of a domain is measured with the average value within the domain on a binarized image with Otsu- binarization algorithm after CLAHE contrast enhancement in ImageJ. The packing efficiency is calculated using the same approach described previously 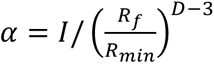, where I is the average chromatin intensity of a domain, R_f_ is the size of the domain, R_min_ is the size of random chromatin polymer chain at which the mass scaling follows the exponential trend *M* ∝ *r*^*D*^ and D is the packing scaling exponent of the domain packing behavior^3^.

The categorization of domains was done by comparison to the median size and packing efficiency of all domains in the control group. The small size and low packing efficiency group is “nascent”, the large size and low packing efficiency group is “decaying”, the high packing efficiency, small size and large size group are “small mature” and “large mature” respectively. After categorization, a detailed radial analysis is done within each group by calculating the mean chromatin intensity as a function of radial distance from the center of the domain.

### PWS Imaging

PWS is a label-free live cell imaging technique that uses the interference in backscattered light from macromolecular structures such as chromatin to survey intracellular structure below the diffraction limit of light. Although individual domains are not directly resolved, PWS nanoscopy quantifies the proportion of chromatin organized into domains, *D*_*nucleus*_, which is proportional to the weighted average of the fractal dimension values of individual domains, 〈*D*_*domain*_〉, and the volume fraction of the packing domains. *D*_*nucleus*_ is also proportional to the fractional moving mass, FMM, defined as the product of the average mass of chromatin clusters that move coherently together per volume fraction of mobile chromatin, and the intranuclear effective diffusion rate of chromatin^33,68,69^. Consequently, increased *D*_*nucleus*_ represents a higher likelihood of chromatin organizing efficiently into domains and vice versa (**Fig. 1c**) (**SI Fig. 1d**). Likewise, increased FMM represents coherent polynucleosome movement, consistent with an increased likelihood of domain formation events. Higher FMM might be due to having a greater volume fraction of mobile vs immobile chromatin or larger chromatin clusters moving as a single unit. The diffusion rate, better understood as the rate of change in chromatin mass-density, can be extracted from the temporal autocorrelation function decay curve by collecting the PWS signal over time at a single wavelength (550 nm). Diffusion provides a useful proxy for measuring the rate of macromolecular motion where smaller chromatin clusters typically correspond to a higher effective diffusion coefficient. A detailed explanation of each PWS imaging modality is below:

#### Dual PWS Imaging

Briefly, PWS measures the spectral interference signal resulting from internal light scattering originating from nuclear chromatin. This is related to variations in the refractive index distribution (Σ) (extracted by calculating the standard deviation of the spectral interference at each pixel), characterized by the chromatin packing scaling (*D*). *D* was calculated using maps of Σ, as previously described^6,68,70^. Measurements were normalized by the reflectance of the glass medium interface (i.e., to an independent reference measurement acquired in a region lacking cells on the dish). This allows us to obtain the interference signal directly related to refractive index (RI) fluctuations within the cell. Although it is a diffraction-limited imaging modality, PWS can measure chromatin density variations because the RI is proportional to the local density of macromolecules (e.g., DNA, RNA, proteins). Therefore, the standard deviation of the RI (Σ) is proportional to nanoscale density variations and can be used to characterize packing scaling behavior of packing domains with length scale sensitivity around 20 – 200 nm, depending on sample thickness and height. Changes in *D* resulting from each condition are quantified by averaging over nearly 2000 cells, taken across 3 technical replicates. Live-cell PWS measurements obtained using a commercial inverted microscope (Leica, DMIRB) using a Hamamatsu Image-EM charge-coupled device (CCD) camera (C9100-13) coupled to a liquid crystal tunable filter (LCTF, CRi VariSpec) to acquire monochromatic, spectrally resolved images ranging from 500-700 nm at 2-nm intervals as previously described^6,68,70^. Broadband illumination is provided by a broad-spectrum white light LED source (Xcite- 120 LED, Excelitas). The system is equipped with a long pass filter (Semrock BLP01- 405R-25) and a 63x oil immersion objective (Leica HCX PL APO). Cells were imaged under physiological conditions (37°C and 5% CO_2_) using a stage top incubator (In vivo Scientific; Stage Top Systems). All cells were given 48 hours to re-adhere before treatment (for treated cells) and imaging.

#### Dynamic PWS Measurements

Dynamic PWS measurements were obtained as previously described^68^. Briefly, dynamics measurements (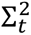, fractional moving mass (*m*_f_), and diffusion) are collected by acquiring multiple backscattered wide-field images at a single wavelength (550 nm) over time (acquisition time), to produce a three-dimensional image cube, where 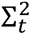 is temporal interference and t is time. Diffusion is extracted by calculating the decay rate of the autocorrelation of the temporal interference. The fractional moving mass is calculated by normalizing the variance of 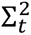 at each pixel. Using the equations and parameters explained in detail in the supplementary information of our recent publication^68^, the fractional moving mass is obtained by using the following equation to normalize 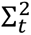 by *ρ*_0_, the density of a typical macromolecular cluster:

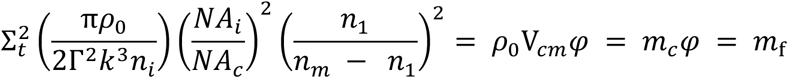

With this normalization, 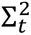 is equivalent to *m*_f_, which measures the mass moving within the sample. This value is calculated from the product of the mass of the typical moving cluster (*m*_*c*_) and the volume fraction of mobile mass (*φ*). *m*_*c*_ is obtained by *m*_*c*_ = V_*cm*_*ρ*_4_, where V_*cm*_is the volume of the typical moving macromolecular cluster. To calculate this normalization, we approximate *n*_*m*_ = 1.43 as the refractive index (RI) of a nucleosome, *n*_1_ = 1.37 as the RI of a nucleus, *n*_*i*_ = 1.518 as the refractive index of the immersion oil, and *ρ*_4_ = 0.55 g *cm*^−3^ as the dry density of a nucleosome. Additionally, *k* = 1.57E5 cm^−1^ is the scalar wavenumber of the illumination light, and Γ is a Fresnel intensity coefficient for normal incidence. *NA*_*c*_ = 1.49 is the numerical aperture (NA) of collection and *NA*_*i*_ = 0.52 is the NA of illumination. As stated in the aformentioned publication, 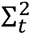 is sensitive to instrument parameters such as the depth of field, substrate refractive index, etc. These dependencies are removed through normalization with the proper pre-factor calculated above for obtaining biological measurements. It should also be noted that backscattered intensity is prone to errors along the transverse direction. Due to these variations, these parameters are more accurate when calculating the expected value over each pixel.

### Protein Detection

HCT116-POLR2A-AID2 cells were lysed using Radio Immuno Precipitation Assay (RIPA) buffer (Sigma-Aldrich, #R0278) with protease inhibitor added (Sigma-Aldrich, #P8340). Cell lysates were quantified with a standard Bradford assay using the Protein Assay Dye Concentrate (BioRad, #500–0006) and BSA as a control. Heat denatured protein samples were resolved on a 4–12% bis–tris gradient gel, transferred to a PVDF membrane using the Life Technologies Invitrogen iBlot Dry Transfer System (Thermo Fisher Scientific, IB1001) (20 V for 7 min), and blocked in 5% nonfat dried milk (BioRad, #120–6404) in 1 × TBST. Whole-cell lysates were blotted against the following primary antibodies: anti-POLR2A (Abcam, ab5408, 1:200 dilution). The following secondary antibodies were used: anti-rabbit IgG HRP (Promega, #W4018). Blots were incubated with the primary antibody overnight at 4 °C, followed by incubation with the secondary antibodies for 1 h at room temperature. To develop blots for protein detection, chemiluminescent substrates were used (Thermo Fischer Scientific, #32106). To quantify the western blot bands, we used the iBright Analysis Software from Thermo Fisher Scientific to define bands as regions of interest. By measuring the mean grey intensity values, the final relative quantification values were calculated as the ratio of each protein band relative to the lane’s loading control for all three replicates. All antibodies used in this study are listed on **Table 3**.

### Flow Cytometry

Flow cytometry analysis for HCT116-POLR2A-AID2 cells to determine proper auxin treatment concentration was performed on a BD LSRFortessa Cell Analyzer FACSymphony S6 SORP system, located at the Robert H. Lurie Comprehensive Cancer Center Flow Cytometry Core Facility at Northwestern University in Evanston, IL. For all FACS analysis, the same protocol was used. After degrading endogenous POLR2A, cells were harvested and analyzed. Briefly, cells were washed with DPBS (Gibco, #14190–144), trypsinized (Gibco, #25200–056), neutralized with media, and then centrifuged at 500 × g for 5 min. Cells were then resuspended in cold FACS buffer (DPBS with 1% of BSA and 2mM EDTA added) at 4°C and immediately analyzed for GFP signal. All flow cytometry data were analyzed using FlowJo software v.10.6.1.

### Coefficient of Variation Analysis

To assess chromatin compaction through the Coefficient of Variation (CoV) analysis, DAPI-stained cells (see section Fixed-cell immunofluorescence) treated with Auxin (see section “Auxin treatment”) were imaged on a Nikon SoRa Spinning Disk confocal microscope (see section “Confocal imaging”). Following a published workflow, we used ImageJ to create masks of each nucleus. The coefficient of variation of individual nuclei was calculated in MATLAB, with CoV = σ/μ, where σ represents the standard deviation of the intensity values and μ representing the mean value of intensity of the nucleus^71^.

### Quantification and Statistical Analysis

Statistical analysis was performed using base functions in R. Pairwise comparisons were calculated on datasets consisting of, at a minimum, three independent replicate samples using two-tailed unpaired t test. Experimental data are presented as the mean ± SEM unless otherwise stated in figure legends. The type of statistical test is specified in each case. For image data, significance was calculated using paired student t test. Multiple comparison corrections were applied for datasets when comparing more than two groups and multiple comparisons. A P value of < 0.05 was considered significant. Statistical significance levels are denoted as follows: ns = not significant; *P < 0.05; **P < 0.01; ***P < 0.001; ****P < 0.0001. Sample numbers (# of nuclei, n), the number of replicates (N), and the type of statistical test used is indicated in figure legends.

## Author contributions

Conceptualization: LMC, LMA, VB, KM

Formal Analysis: RG, NA, WSL, NP, KW

Methodology: LMC, LMA, RG, NA, WSL, SH, MK

Investigation: LMC, LMA, EP, KLM, WSL, SH, NP, KW, TK

Visualization: LMC

Funding acquisition: VB, LMA, KM

Project administration: LMA, VB

Supervision: LMA, VB

Writing: LMC, LMA

## Acknowledgements

Philanthropic support was generously received from Rob and Kristin Goldman, the Christina Carinato Charitable Foundation, Mark E. Holliday and Mrs. Ingeborg Schneider, and Mr. David Sachs. This research was supported in part through the computational resources and staff contributions provided for by the Quest high performance computing facility at Northwestern University which is jointly supported by the Office of the Provost, the Office for Research, and Northwestern University Information Technology. This research was supported in part through the computational resources and staff contributions provided by the Genomics Compute Cluster which is jointly supported by the Feinberg School of Medicine, the Center for Genetic Medicine, and Feinberg’s Department of Biochemistry and Molecular Genetics, the Office of the Provost, the Office for Research, and Northwestern Information Technology. The Genomics Compute Cluster is part of Quest, Northwestern University’s high performance computing facility, with the purpose of advancing research in genomics. We appreciate the generous support from the ENCODE Consortium in the generation and dissemination of publicly available datasets. We specifically want to thank the labs of John Lis, Bas van Steensel, Bradley Bernstein, and Richard Myers for the generation and publication of the data utilized within this manuscript. We also want to thank Shelby Blythe, Yue Yang, and Xiaozhong Wang at Northwestern University for their guidance on this project.

## Funding

National Institutes of Health grant U54 CA268084

National Institutes of Health grant R01CA228272

National Institutes of Health grant U54 CA261694

NIH Training Grant NIH: T32 GM142604

NIH Training Grant T32AI083216 (LMA) NSF: EFMA-1830961

The Center for Physical Genomics and Engineering at Northwestern University

CURE Childhood Cancer Early Investigator Grant

Alex’s Lemonade Stand Foundation ‘A’ award grant #23-28271

Pediatric Cancer Research Foundation

## Competing interests

Authors declare that they have no competing interests related to this work.

## Data and materials availability

All of the data, code used for analysis, and relevant materials have been uploaded to the following locations. Publicly available Micro-C and Pro-Seq data were obtained from the ENCODE Consortium, GEO, or 4DN with the relevant accession numbers and file names are shared in the supplementary information in **Table 5**. Code used for analysis and figure has been uploaded to GitHub at https://github.com/BackmanLab/Transcription. Processed data along with all code needed to generate figures is available at https://datadryad.org/stash/share/wGKlTXjBvL9uBW8HgMOMgermP0dki79WXs25JnSZxM4. All sequencing data has been deposited at GEO under the following accession numbers - RNAseq: GSE279127 , HiC: GSE279166.

## Notes

### Competing Interest Statement

The authors have declared no competing interest.

