## Supplemental Information for "Gene transcription and chromatin packing domains form a self- organizing system"

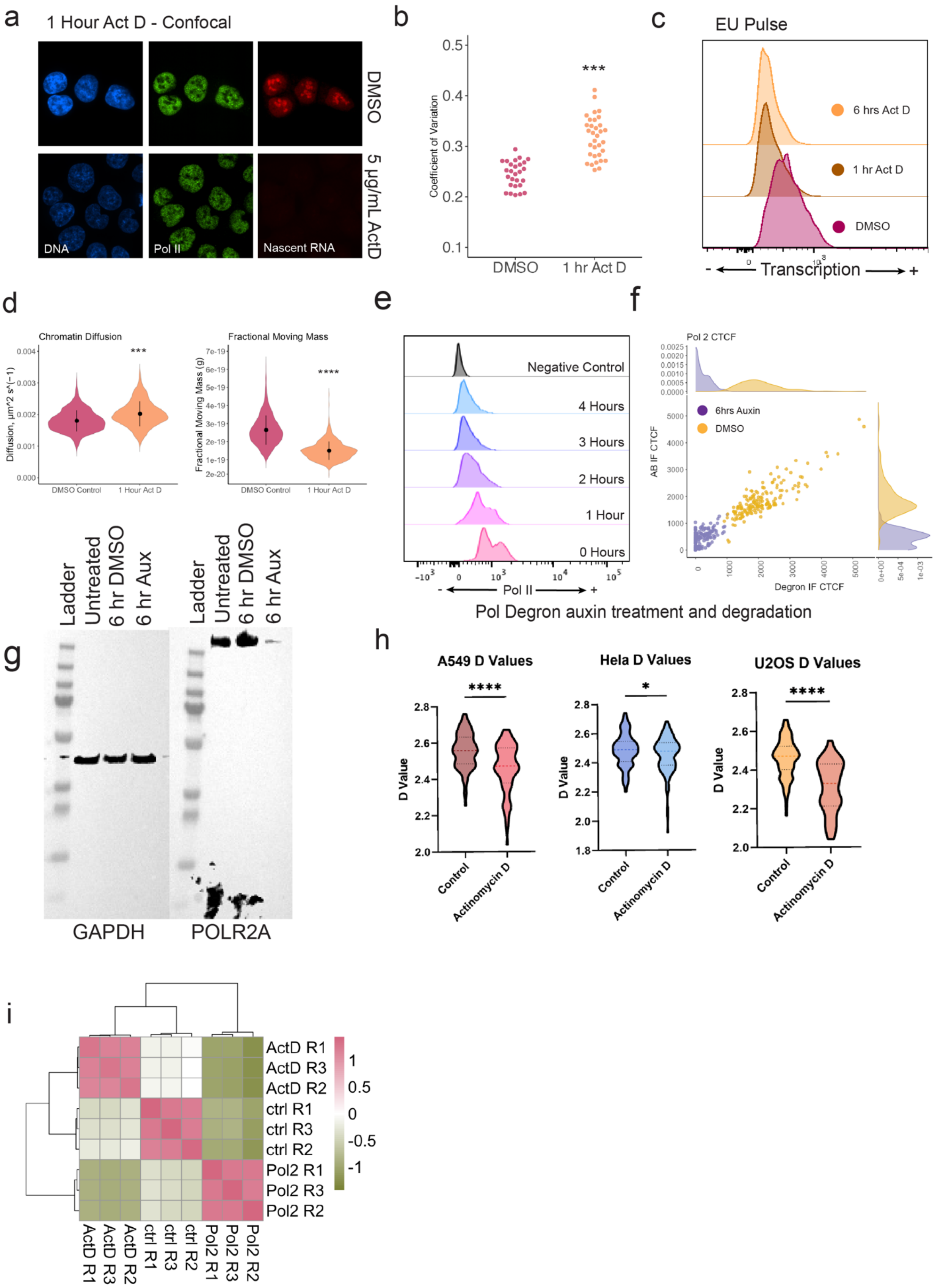
a
1 Hour Act D - Confocal
DMSO
5 μg/mL ActD
DNA
Pol II
Nascent RNA
b
Coefficient of Variation
***
DMSO
1 hr Act D
c
EU Pulse
6 hrs Act D
1 hr Act D
DMSO
- Transcription +
d
Chromatin Diffusion
Diffusion, μm^2 s^(−1)
***
DMSO Control
1 Hour Act D
Fractional Moving Mass
Fractional Moving Mass (g)
****
e
Negative Control
4 Hours
3 Hours
2 Hours
1 Hour
0 Hours
- Pol II +
Pol Degron auxin treatment and degradation
f
Pol 2 CTCF
6hrs Auxin
DMSO
AB IF CTCF
Degron IF CTCF
g
Ladder
Untreated
6 hr DMSO
6 hr Aux
GAPDH
POLR2A
h
A549 D Values
Hela D Values
U2OS D Values
D Value
Control
Actinomycin D
****
*
i
ActD R1
ActD R3
ActD R2
ctrl R1
ctrl R3
ctrl R2
Pol2 R1
Pol2 R3
Pol2 R2

**SI Figure 1.** Treatment with Actinomycin D (5 μg/mL) for 1 hour leads to chromatin compaction and abrogation of transcription and validation of HCT116 POLR2A-AID2 line. (**A**) widefield fluorescent microscopy of ActD treated cells or DMSO control cells: DAPI stained DNA (blue), labelled POLR2A (green), EU-labelled nascent RNA (red). (**B**) Coefficient of variation analysis for DAPI stained DNA before and after treatment. The coefficient of variation of individual nuclei was calculated in MATLAB, with CoV = σ/μ, where σ represents the standard deviation of the intensity values and μ representing the mean value of intensity of the nucleus. See methods for more details. (**C**) Flow cytometry analysis of EU-labelled nascent RNA before treatment and at 1 or 6 hours of treatment, showing loss of RNA signal after treatment. (**D**) PWS following treatment: average diffusion and fractional moving mass of chromatin domain nuclear average, respectively. Data was compiled from three independent biological replicates. (DMSO: N = 475; ActD: N=541). (**E**) Flow cytometry analysis of target protein GFP signal following treatment with auxin shows depletion of tagged POLR2A by 4 hours. (**F**) Verification of POLR2A depletion by quantification of degron GFP signal and immunofluorescent signal showing over 90% of the population is without detectable protein and degron-tagged protein signal is consistent with total protein signal. Axes represent corrected-total-cell-fluorescence in relative units. Axes represent corrected-total-cell-fluorescence in relative units. (**G**) Western blot of POLR2A-AID2 line after 6 hours of degradation shows full degradation of POLR2A protein. Western blot was completed once, to confirm earlier WB validation prior to our acquisition of this line. (**H**) Nuclear scaling analysis of A549, HeLa, and U2OS cells following treatment using PWS. (**I**) Sample Pearson correlation between replicates for RNA-seq. For **B** and **D**, significance was calculated by unpaired *t* test with Welch's correction applied (**** < 0.0001).

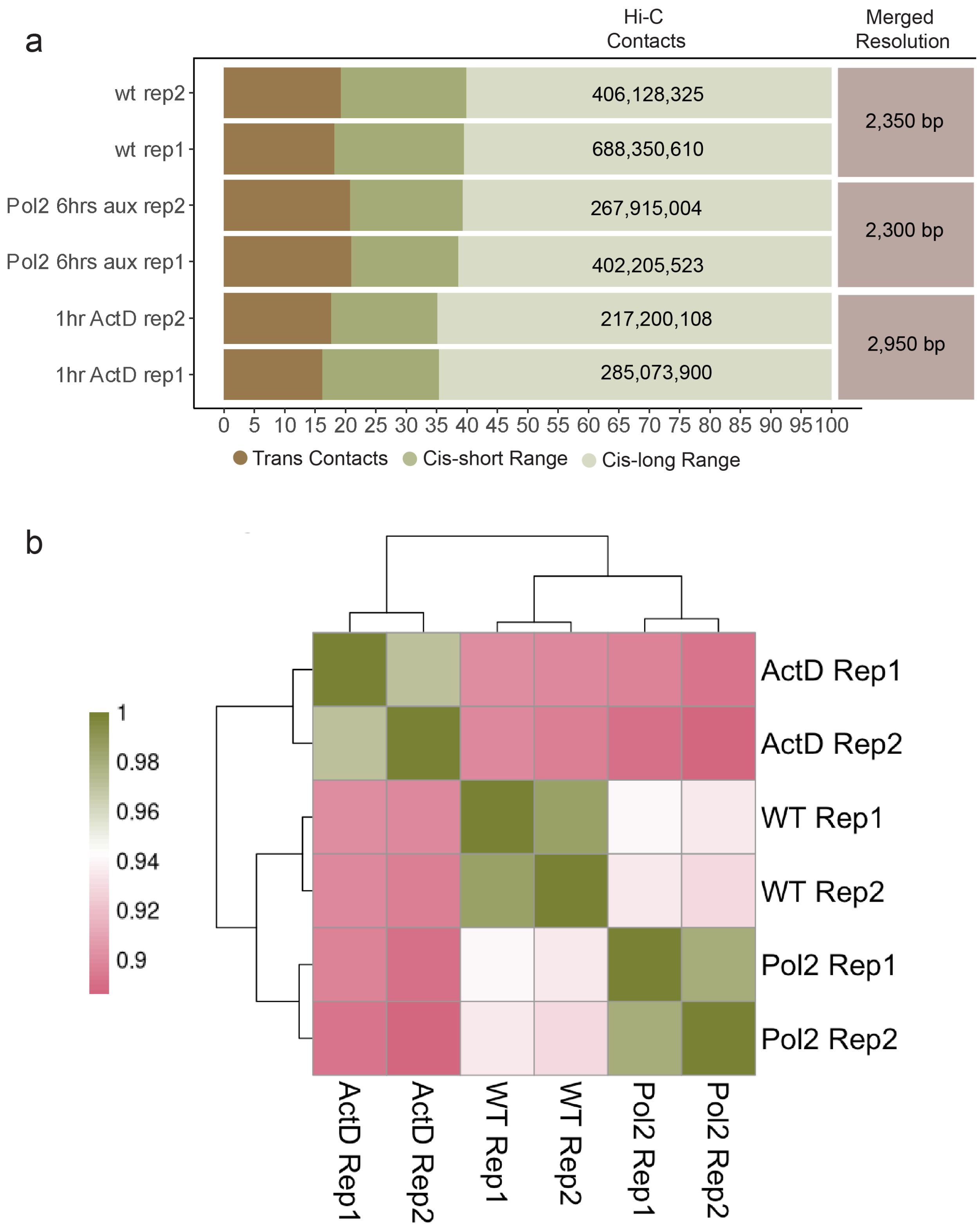
a
Hi-C Contacts
Merged Resolution
wt rep2
wt rep1
Pol2 6hrs aux rep2
Pol2 6hrs aux rep1
1hr ActD rep2
1hr ActD rep1
406,128,325
688,350,610
267,915,004
402,205,523
217,200,108
285,073,900
2,350 bp
2,300 bp
2,950 bp
0 5 10 15 20 25 30 35 40 45 50 55 60 65 70 75 80 85 90 95 100
Trans Contacts
Cis-short Range
Cis-long Range
b
1
0.98
0.96
0.94
0.92
0.9
ActD Rep1
ActD Rep2
WT Rep1
WT Rep2
Pol2 Rep1
Pol2 Rep2

**SI Figure 2.** Two technical Hi-C replicates (treated in separate dishes on the same day) each were generated from POLR2A-AID2 degron line treated with auxin for 6 hours, DMSO for 6 hours, or Actinomycin D (5 µg/mL) for 1 hour. (**A**) Total number of contacts, trans contacts, and cis contacts for each replicate; resolution of merged replicates. Y axis = % of total contacts. Hi-C contact number is the total number of contacts in library calculated by Juicer pipeline after removal of ambiguous contacts and duplicates. (**B**) The stratum adjusted correlation coefficient (SCC) generated for all replicates shows high correlation between technical replicates.

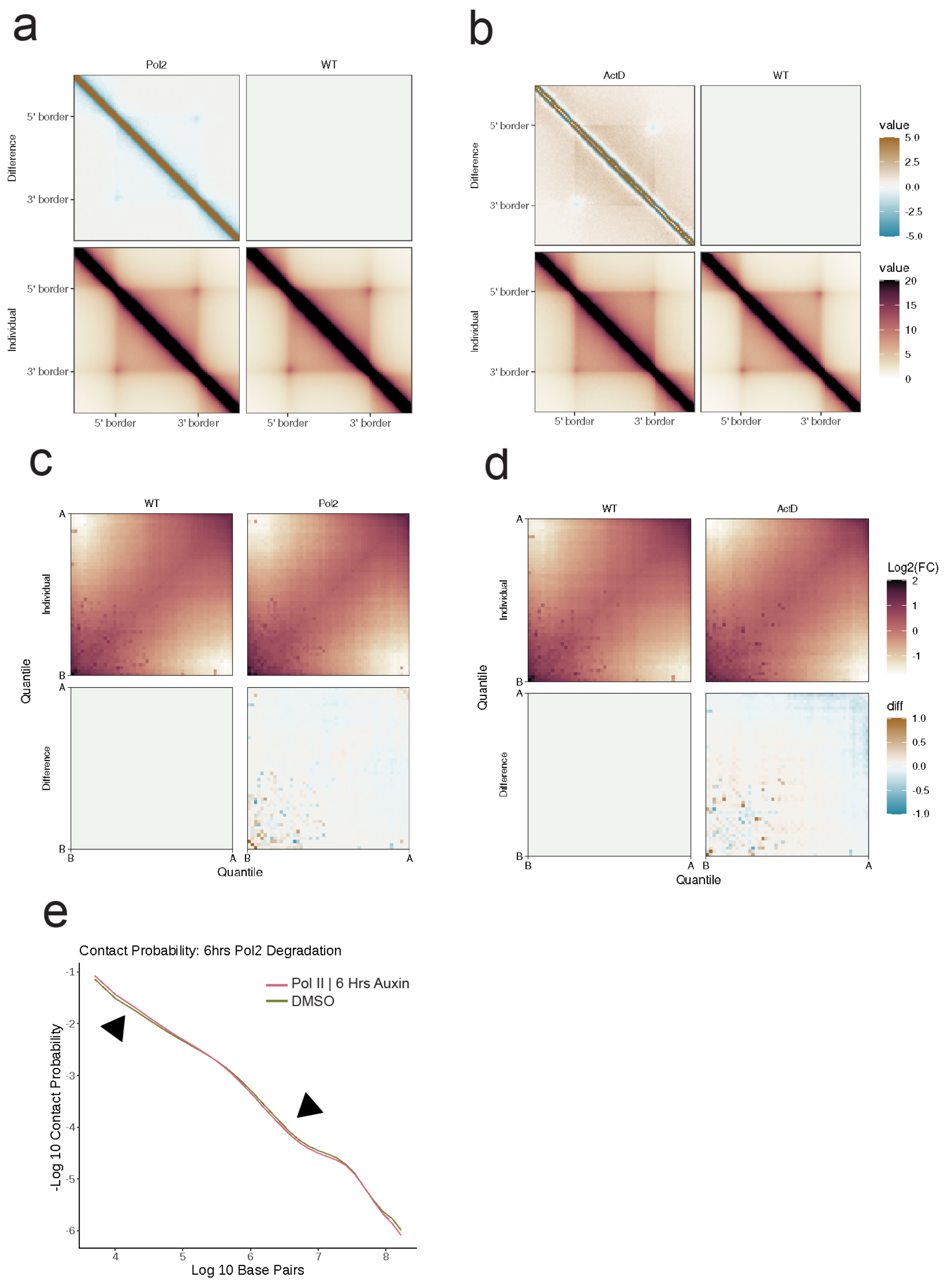


**SI Figure 3.** Additional Hi-C analysis. POLR2A-AID2 degron line was treated with auxin for 6 hours, DMSO for 6 hours, or Actinomycin D (5 µg/mL) for 1 hour prior to contact mapping. For all conditions: (**A-B**) TAD insulation plots showing no change to TADs between conditions, (**C-D**) compartment pile-up plot showing no change to compartments (both TAD and compartment pileup plots were generated using GENOVA. See methods for more details), and (**E**) Relative contact probability.

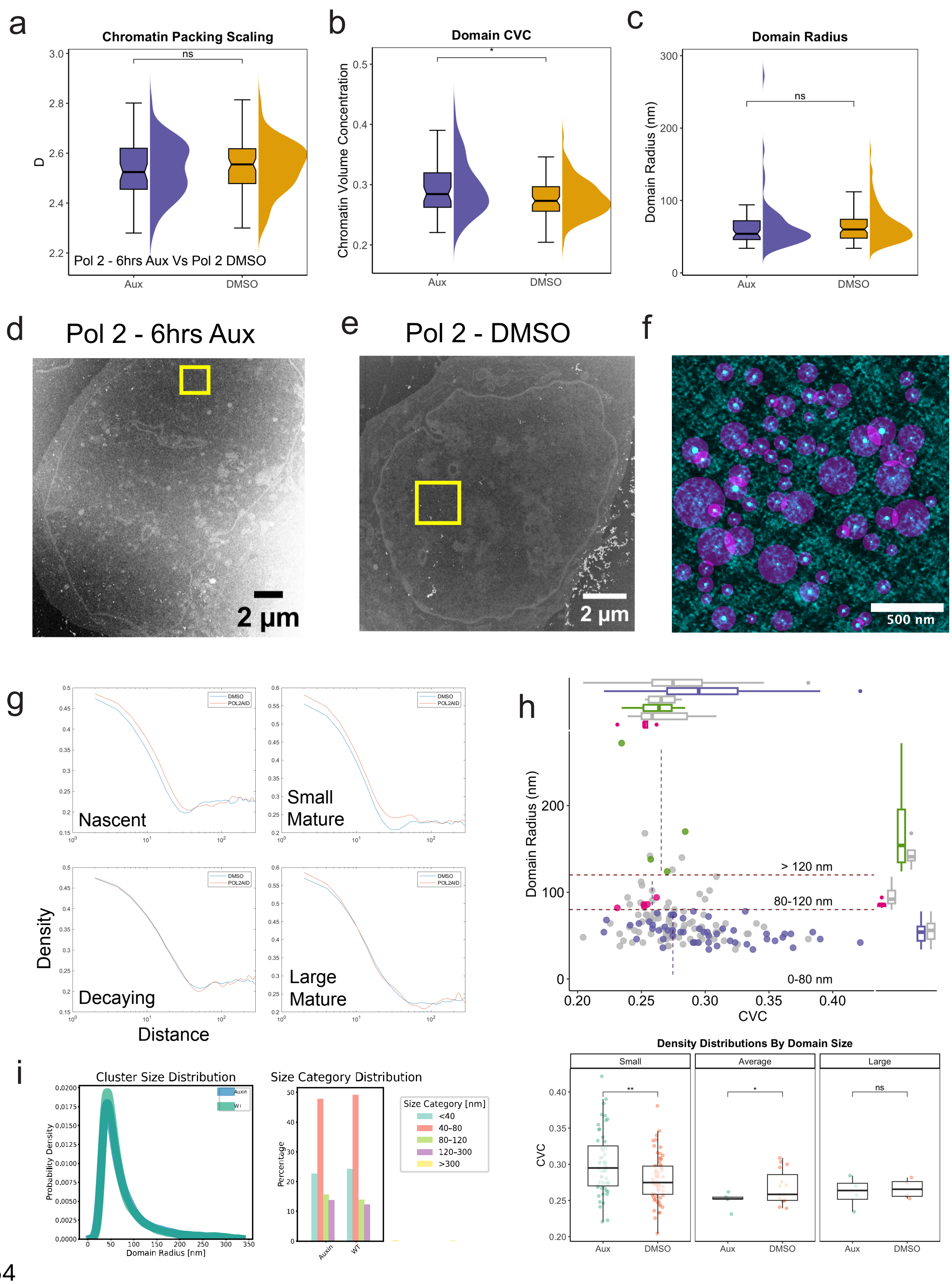
a
Chromatin Packing Scaling
ns
D
Pol 2 - 6hrs Aux Vs Pol 2 DMSO
Aux
DMSO
b
Domain CVC
*
Chromatin Volume Concentration
Aux
DMSO
c
Domain Radius
ns
Domain Radius (nm)
Aux
DMSO
d
Pol 2 - 6hrs Aux
2 μm
e
Pol 2 - DMSO
2 μm
f
500 nm
g
Nascent
Small Mature
Decaying
Large Mature
Density
Distance
DMSO
POL2AID
h
Domain Radius (nm)
> 120 nm
80-120 nm
0-80 nm
CVC
Density Distributions By Domain Size
Small
Average
Large
**
*
ns
CVC
Aux
DMSO
i
Cluster Size Distribution
Probability Density
Domain Radius [nm]
Auxin
WT
Size Category Distribution
Percentage
Size Category [nm]
<40
40–80
80–120
120–300
>300
Auxin
WT

**SI Figure 4.** Supplementary ChromSTEM analysis. (**A-C**) ChromSTEM following treatment: packing scaling, chromatin volume concentration, and domain radius of individual domains, respectively (Auxin: N = 72; DMSO: N=61). Significance was calculated by unpaired *t* test with Welch’s correction applied (**** < 0.0001). (**D-E**) Scanning Transmission Electron Microscope HAADF image of nucleus. The box shows the region where ChromSTEM tomography data is collected. Scale bar = 200nm. (**F**) Example ChromSTEM tomogram overlayed with identified domains center (highlighted in blue) based on local maxima (ImageJ) and domain size in purple used for **Figure 4f**. (**G**) Radial Density (y-axis) vs Distance from domain center (x-axis) average within domain groups. (**H**) Domain radius versus chromatin volume concentration (density) for all domains on ChromSTEM. Second plot bins domains by size and compares chromatin density. (**I**)  Size distribution of H3K9me3 cores. Significance was calculated by unpaired *t* test with Welch’s correction applied (**** < 0.0001).

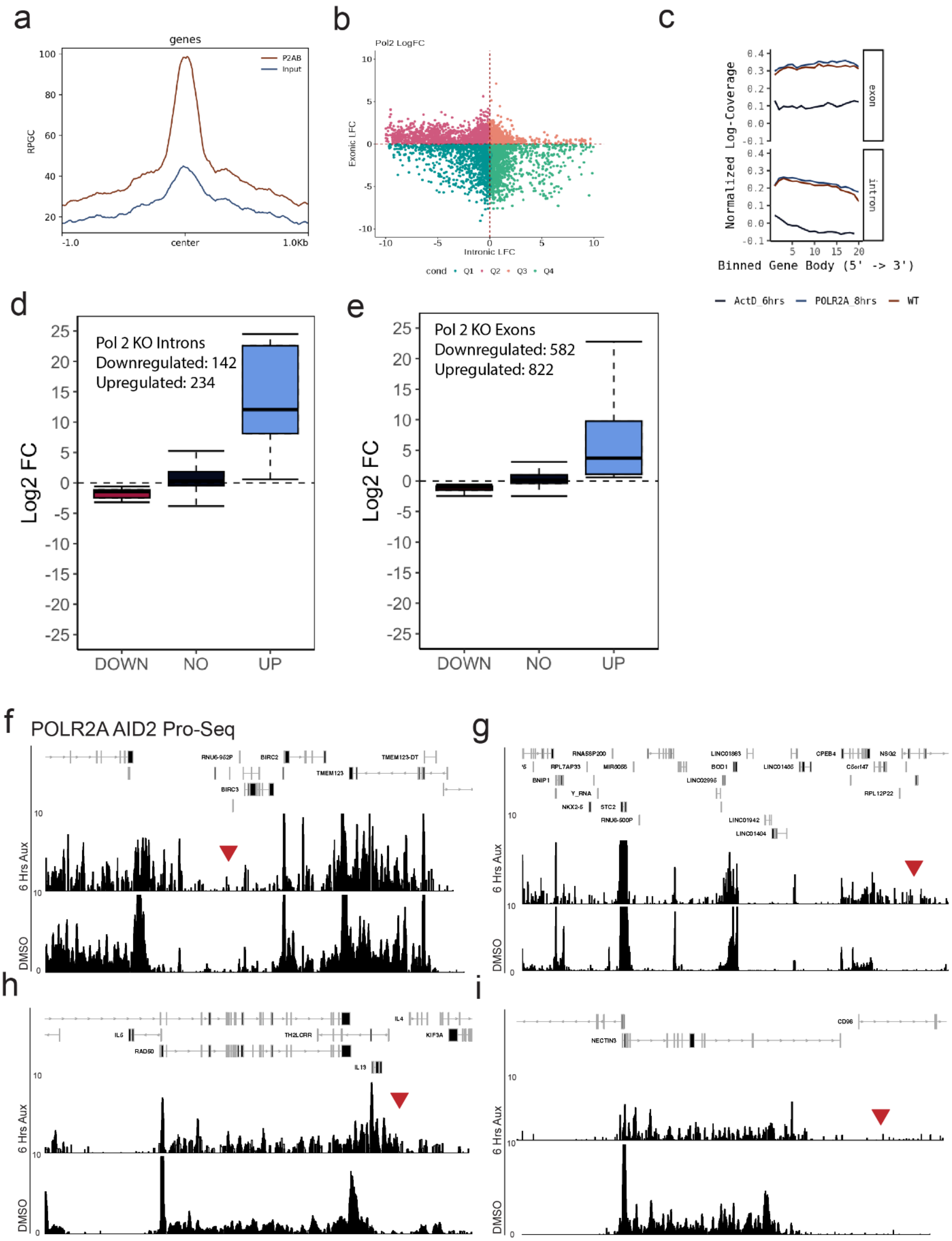
a
genes
P2AB
Input
RPGC
-1.0
center
1.0Kb
b
Pol2 LogFC
Exonic LFC
Intronic LFC
cond
Q1
Q2
Q3
Q4
c
Normalized Log-Coverage
exon
intron
Binned Gene Body (5' -> 3')
ActD_6hrs
POLR2A_8hrs
WT
d
Pol 2 KO Introns
Downregulated: 142
Upregulated: 234
Log2 FC
DOWN
NO
UP
e
Pol 2 KO Exons
Downregulated: 582
Upregulated: 822
Log2 FC
DOWN
NO
UP
f
POLR2A AID2 Pro-Seq
6 Hrs Aux
DMSO
g
6 Hrs Aux
DMSO
h
6 Hrs Aux
DMSO
i
6 Hrs Aux
DMSO

**SI Figure 5.** Supplementary transcriptomic analysis. (**A**) Pileup plot of BPM normalized signal from POLR2A RIP-seq peaks in WT HCT116 cells. (**B**) Exon log fold change and intron log fold change generated using Index package plotted for each gene. (**C**) Nascent intron and exon signal from POLR2A-degraded cells was averaged across 20 bins for all gene bodies. Coverage was normalized, log2 transformed, and plotted. ActD shows loss of intronic signal. (**D**) and (**E**) Loss of Pol-II upregulates intronic and exonic nascent transcripts within gene bodies. Analysis of nascent expression in EU-seq generated from POLR2A-AID2 degron line treated with auxin or DMSO for 8 hours or ActD (5 μg/mL) for 6 hours. Differentially expressed introns and exons in POLR2A-depleted cells ($P_{adj} < 0.05$, LFC > abs(1) ). (**F-I**) Publicly available Pro-Seq featuring POLR2A-degraded cells. Genomic loci correspond to **Figure 6D-G** and red arrows point to same 5' read-through regions.

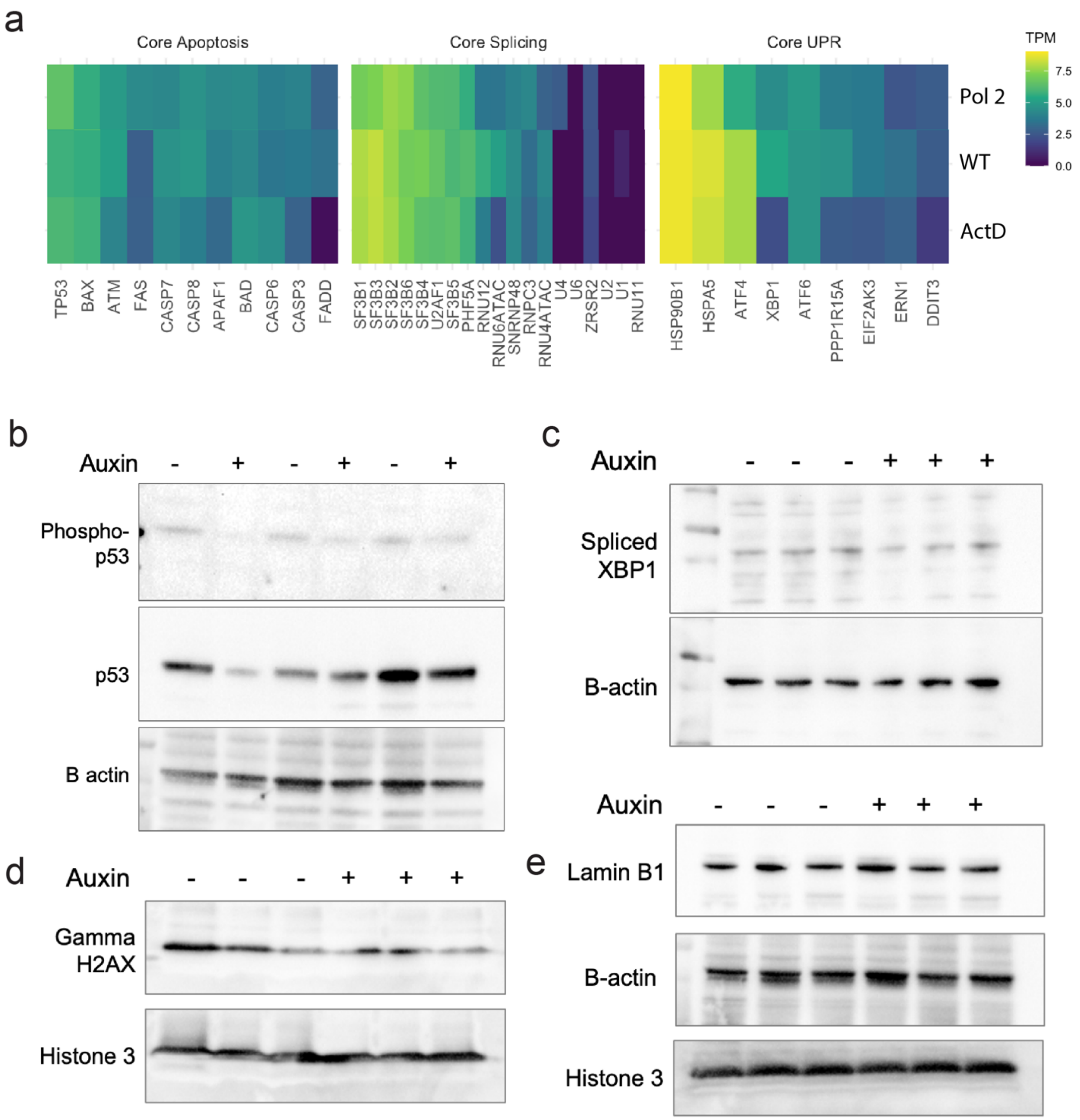


**SI Figure 6.** Analysis of normalized gene counts in GO biological process terms and western blots of corresponding markers (**A**) TPM normalized counts for core splicing, core apoptosis, and core UPR genes in each condition. Core lists were taken from GO biological processes lists. (**B-E**) Western blots of auxin treated POLR2A-AID2 cells or DMSO treated POLR2A-AID2 cells (6 hours): (**B**) P53 and Phosphorylated P53, (**C**) s-XBP1, (**D**) Gamma H2AX, and (**E**) Lamin B1.
